# *WDR88*, *CCDC11*, and *ARPP21* genes indulge profoundly in the desmoplastic retort to prostate and breast cancer metastasis

**DOI:** 10.1101/178566

**Authors:** Rajni Jaiswal, Shaurya Jauhari, S.A.M. Rizvi

## Abstract

Microarray technology has unlocked doors to a multitude of open analysis problems that if conceived with efficacy may uncover varied genotypic and phenotypic traits. Algorithms belonging to different cultures in computer science have been applied to gene expression data to derive correlation and stratification parameters. While most outcomes are subject to clinical validation, majority of which get declined, the search for the precisely targeted therapeutic agents is still on. This paper is an effort in the similar direction and strives to delineate genes with significant stromal signatures. We suggest a corroborative indulgence of a human laterality disorder gene, CCDC11 in the metastasis, in addition to the role of *WDR88* and *ARPP21* genes has been further materialized in the analysis. Another standout aspect of the study has been the associated implications of the genes in rare disorders of male breast and female prostate cancers. There is also a threshold proposal that stratifies “safe” expression space for genes. Complimentarily, the manuscript serves as an expedient protocol for anyone seeking microarray data analysis, particularly in R.

## 1. Introduction

There appears nothing proverbially eerie about the technology at the get go. Microarrays usage throve with (Schena et al. 1995) and were originally applied to harbour global gene expression (DeRisi et al. 1997; DeRisi et al. 1996) in association with yeast studies. With the proliferation of data pertaining to medication and that too in the digital proforma, it is crucial to constantly challenge and update the current configuration of systems that are being used to analyse it [genomic data] for compliance to the medical care. NGS is one such advancement that was gullible to the geneticists. Unlike the microarray data that catalogues gene expression values under a predefined probe, the RNA-seq data from NGS documents expression range in totality (Uziela & Honkela 2013). RNA sequencing technology pictures a comprehensive view of the transcriptome with the data being reproducible for novel discoveries yielded by disparate analyses. RNA sequencing is also helpful in detection of structural variations as gene fusions, alternative splicing events, etc. But microarrays still continue to provide a relatively affordable *first-foot* to genomics, bearing robustness and short turn-around time. With significant disparities owing to the definite and specific backgrounds of the individuals, the genomic data available via microarray format has shown likewise results when particular maladies come into question as cancer, diabetes, amongst others. The big question however is that could the genes be standardized via ontology driven mechanism so that specific drug targets be known and hunted for. Scientists are always looking for particular biomarkers that can be universally acclaimed and acknowledged. In the current paper, we attempt to underline key players that are actively responsible for representing the metastatic behaviour and proliferation of oncogenic state in a body induced with breast-type and pros-tate-type cancers, in cognizance to stromal reaction. The results are based on a comparative meta-analysis.

Reactive stroma is a response to the aberration into the tissues due to tumor invasion (Planche et al. 2011). Synonymous to desmoplasia, it has also been recognized that the stromal response is exclusive to tumor type. It can be perceived that desmoplastic response is in tandem to carcinogenesis and subsequent metastasis. Thus, it is unstated that desmoplastic reaction is also a prospective antecedent of premalignant stage, as the growth of connective, fibrous tissues around the tumor cells commences. Genetic irregularity in the cells compartmentalized in epithelium represents carcinoma in situ and the lesions initiate cell fibroblasts as a tackling measure. Functionalities of stromal initiation include homeostasis and tissue structure restoration. Chronicled is also that cell division govern mechanism is hampered because of the tumor induction and eventual progression. The amount of reactive stroma is proportional to the disease state (Martin & Rowley 2013). Once the tumor foray infiltrates through the ECM into adjoining host tissues, they become potent to further metastasize. Vascular structures, blood and lymph vessels, ECM, and fibroblasts constitute the stroma (Casey et al. 2009). Diverse studies by (Tuxhorn et al. 2002), (Ayala et al. 2003), (Roepman et al. 2006), and (Finak et al. 2006) implemented LCM to scrutinize gene expression profiles of tumor stroma (breast) versus normal epithelium and clinched that the alterations in the stromal microenvironment is comparative to the tumor progression.

In the following work, we attempt to ascertain genes that are prominent to tumor progression and subsequent stromal response. This may aid identification of key pathways (genes instituted) that are liable for the cancer metastasis. As the dataset may reveal, we attempt to analyse breast and prostate oncogenes.

This paper is organized as follows:

First, the developments in the breast cancer and prostate research, over the years, are catalogued. Various data analysis methodologies that have inferred some very seminal results have been underlined. We then present our viewpoint and improvements in the domain and propose a novel algorithm to analyse cancer stroma data. As it would necessitate, the sifted targets are subjected to validation; but due to accessibility constraints, could only be done via available erstwhile published research work. Their [genes] analysis can further substantiate our studies for preventing the spread of cancer to the other tissues through pathway blockage and rendering them benign through a drug treatment.

The statistical analysis and visualizations are covered with R language (version 3.2.3) (interface used is RStudio version 0.99.491) on a desktop computer with 8 Gb RAM and an Intel i5 CPU with 3.50 Ghz clock speed. For further distillations, MeV version 4.9.0 (Anon n.d.), and Cytoscape are employed.

### 1.1 An Alarming Statistic

A not so long ago article (Kamath et al. 2013), reports that India is overwhelmed with 2.5 million cancer patients in aggregate and close to a million such augmented annually. To put things into perspective, there were 1.7 million and 11.4 million cancer incidences in the South East Asia region and world over, respectively in 2004. According to Globocan data (International Agency for Research on Cancer), India tops the chart with 1.85 million years of healthy life lost due to breast cancer alone. The aftermaths of this malady are equally likely for the rest of the world too.

* *Healthy life lost is defined by years lost owing to premature death and deterioration of health standards on account of a disease induction into the body*.

An elucidation from an erstwhile research confirms that after cervical cancer, breast cancer is highly promulgated amongst Indian women. Also shown is that Indian women are likely to inhibit breast cancer, a decade earlier than their Western counterparts. The paucity of early detection and incompetent control mechanism can largely explain the succumbing rate. Exorbitance in breast cancer cases throughout developing nations is proportionate to varying lifestyle being is unregulated and sporadic, expectancy and delivery of fewer children, and hormonal intervention exemplified by post-menopausal hormonal therapy. The symptoms are profound at a later stage of the malignancy and hence pose greater challenge to review the disease at the initiation. The authors of the study (Kamath et al. 2013) stressed upon the need to exorcise this “ticking time bomb” and called for apt administrative measures for the same.

Prostate cancer, mostly occurring in elderly men, has similar danger trail and accounts for second largest cancer causing deaths in U.S. males after lung cancer (Siegel et al. 2016; Gaylis et al. 2016).

### 1.2 Provenance

When it comes to being most defiant and stubborn, and not to mention “incurable”, cancer is christened far and wide for being the malady that poses serious threat to the manhood. Many of the responsible genes involved in the pathways oriented to oncological disorders have complex and overlapped functioning. Not to mention, some genes remain dormant at an instance and are activated by a particular range of expression level of other corresponding gene[s]. They also tend to become chemotherapy resistant through a self-regulatory mechanism. These attributes account for a thorough and complacent inspection of the various parameters involved in gene functioning, mapped and homed-in.

In the exploration of gene expression data, the magnitude of tissue samples is lower with respect to number of genes that may inevitably lead to overfitting of data and inappropriate results (Shen et al. 2007). Gene selection is critical to elucidate tissue classification as well as to model complex genetic and molecular underpinnings, which explain the relation between genes and varied biological phenomenon. The stability of the analysis model can be accomplished through it.

BRCA1 and BRCA2 are vehemently recognized for hereditary breast and ovarian cancer proneness. These are human genes that produce tumor suppressor proteins that implicitly initiate DNA repair mechanism. If a mutation is detected in any of the aforementioned genes, the susceptibility to espousing tumor inception is high (Anon n.d.). BRCA mutation may lead to the following probabilities:

- *40%-80% for breast cancer*
- 11%-40% for ovarian cancer
- 1%-10% for male breast cancer
- *Up to 39% for prostate cancer*
- 1%-7% for pancreatic cancer

Likewise, if any other relative cancer genes could be deciphered by comprehensively analyzing the gene expression data and establishing their helm in metabolism via clinical validation, we can get closer to disease treatment and increased understanding towards biology. (Bosdet et al. 2013) take BRCA mutation testing to a whole new level by incorporating the Second Generation Sequencing and Third Generation Sequencing procedures, collectively known as NGS, to deal with increasing number of tests that the people are willing to take to judge their cancer proneness. This era of NGS renders reduced cost, greater efficiency and high throughput. The assay defined uses automated small amplicon PCR followed by sample pooling and sequencing with a second-generation instrument.

### 1.3 Androgen Receptor: An observable *cause commune*

Classically abnormality in males associated with prostate cancer, androgen receptor response has been apropos (Yu et al. 2000). AR gene isn’t solely responsible to harbor design and characteristic instructions for sex drive and hair growth, but also facilitate sexual physiognomies. Positioned on the long (q) wing of the X chromosome at the 12^th^ position, the AR gene encompasses cohorts of CAG repeat regions (*triplets* or *trinucleotide repeats*). The strength of quantifiable occurrences of these DNA segments account for the proneness of the prostate cancer and breast cancer; while some studies hold more repeats liable, others blame lesser ones (Yu et al. 2000). Research also depicts that mutations in the AR gene are accountable for prostate cancer instantiation (Nelson 2002) (Giovannucci et al. 1997), albeit somatic in nature. In women, longer CAG repeats and polymorphisms may increase the risk of endometrial and breast cancers (Mehdipour, Pirouzpanah, Kheirollahi, & Atri, 2010).

### 1.4 Gene Selection

While holding candescence to the fact that intergenic regions relegated as “junk DNA” have long been undermined, numerous follow up studies have unraveled that non-coding RNAs, amongst other “dark” regions have a profound effect on regulation of gene expression (Birney et al. 2007) (Carninci et al. 2005) (Cheng et al. 2005) (He et al. 2008). Since microarrays are designed to study gene measurements, the aforementioned parameters are left diluted. This aspect holds its vitality and is sure to influence the end result. Notwithstanding, it has been known that Particle Swarm Optimization Technique (PSO) has been meticulously significant in harnessing gene selection (Shen et al. 2007) (Yuan & Chu 2007) (Shen et al. 2008) (Chuang et al. 2008) (Lin et al. 2008). Other approaches include Artificial Neural Networks (ANN) and Fisher Discriminant Analysis (FDA), to name a few. An ensemble methodology involving Particle Swarm Optimization (PSO) and Support Vector Machines (SVM) has been observed to be particularly critical to feature selection and cornering genes of interest (Yeung et al. 2009).

### 1.5 Elucidation of Cancer Subtypes

Breast cancer is a neoplasm that with distinct subtypes has differently representable histopathological features and response to systemic therapies (Dai et al. 2016). Patient age, tumor size, and axillary lymph node status have been deciding factors as well (Schnitt 2010). Immunohistochemistry (IHC) biomarkers have been classically deployed to ascertain subtyping. They entail Estrogen Receptor (ER), Progesterone Receptor (PR), Androgen Receptor (AR), and Human Epidermal growth factor Receptor 2 (HER2). Back in the 70’s, there were two subtypes that became known to us, viz. (luminal epithelial) ER+ and ER-(Perou et al. 2000) (Sorlie et al. 2003) (Alexe et al. 2007). Triple negative breast cancer is characterized by a cancer subtype devoid of ER, PR, and HER2 gene expressions. Compounds like tamoxifen (for ER), and trastuzumab (for HER2), are tactless in dealing with triple-negative breast cancer. It is chemotherapeutically challenging as it warrants a grouping of disparately rated drugs to target each of the receptor. Owing to its profile, triple negative breast cancer is revered as a basal-type. Another recent study (Vici et al. 2015), illumes reasonableness of the triple positive breast cancer.

From prostate cancer viewpoint, gene fusions between TMPRSS2 and ETS hierarchies have been stressfully documented (Tomlins et al. 2006), and also with ERG genealogy (Penney et al. 2016). Expression levels of genes *MUC1* and *AZGP1* were also shown to categorically underline exclusive subtypes of prostate cancer from clinicopathological stance (Lapointe et al. 2004).

(Herschkowitz et al. 2007) orchestrated a pioneering work that led to elucidation of a novel sub-type pertaining to breast cancer disorder. This new subtype, referred to as Claudin-Low was implicit of low expression genes. Also, traditionally, tumor types could be classified as basal epithelial-like group (ERs), an *ERBB2*-overexpressing group, and a normal breast-like group (Davidson & Liu. 2010). Another feature discovery from a study by (Sorlie et al. 2001) had confirmed the possible subdivision of the ER+ tumor type into two clusters with distinctive gene sets having particular corresponding clinical outcome.

## 2. Results and Discussion

The exegesis is premeditated so as to elucidate a quantifiable threshold that stratifies gene expression space in conjunction to normal and cancer stromal response states. We deliberate to identify key transcriptional features that determine the high dimensional feature space and visualize their inter-linkups via a regulatory network illustration. This is always complimentary to ascertain our knowledge about genes and their pathway-occurrence motifs… (Figure 1)

**Figure 1.**
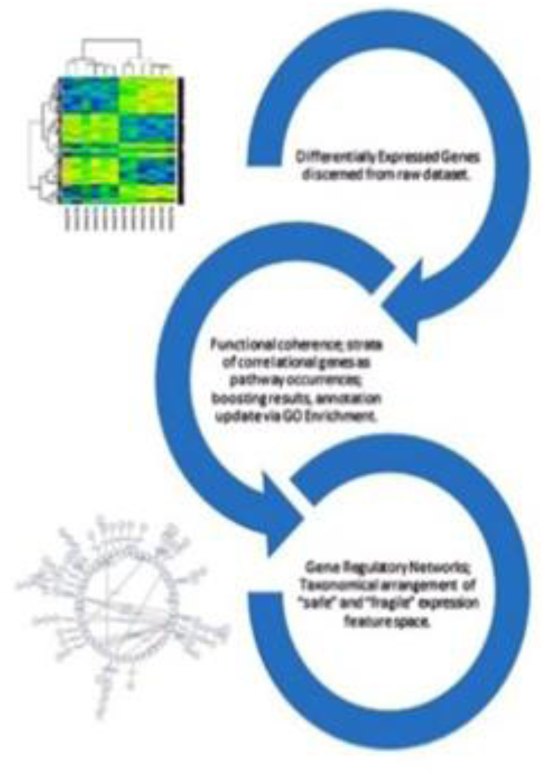
Illustration of workflow.

### 2.1 Anonymous Genes/ Probes

We identified 27236 entries, while scrutinizing the annotation fields of the dataset that eluded ontological reference. There are also considerable amount of genes whose expression values are catalogued under incongruent probes, resulting in their multiple incidences. This is a purported case of genes’ splice variants, as the dataset suggests. While we aim to identify DEG and construct a respective GRN, there is also a prudence of elucidating functionally coherent genes that may unravel profiling of all or few anonymous genes. Thus, it appears duteous to abandon blank values to sustain quality of biological interpretation and germaneness.

### 2.2 Normality and Data stabilization

The data appears normalized data sans log transformation. Hence it is log-transformed and metamorphosed to render mean=0, and standard deviation=1, i.e. it followed normal distribution. Since, the normality isn’t skewed as a result of multiple comparisons problem (Dunn 1961), as we’re not envisioning multitude of significance values, there is no proliferation of Type I error occurrence anomalies… (Figure 2)

**Figure 2.**
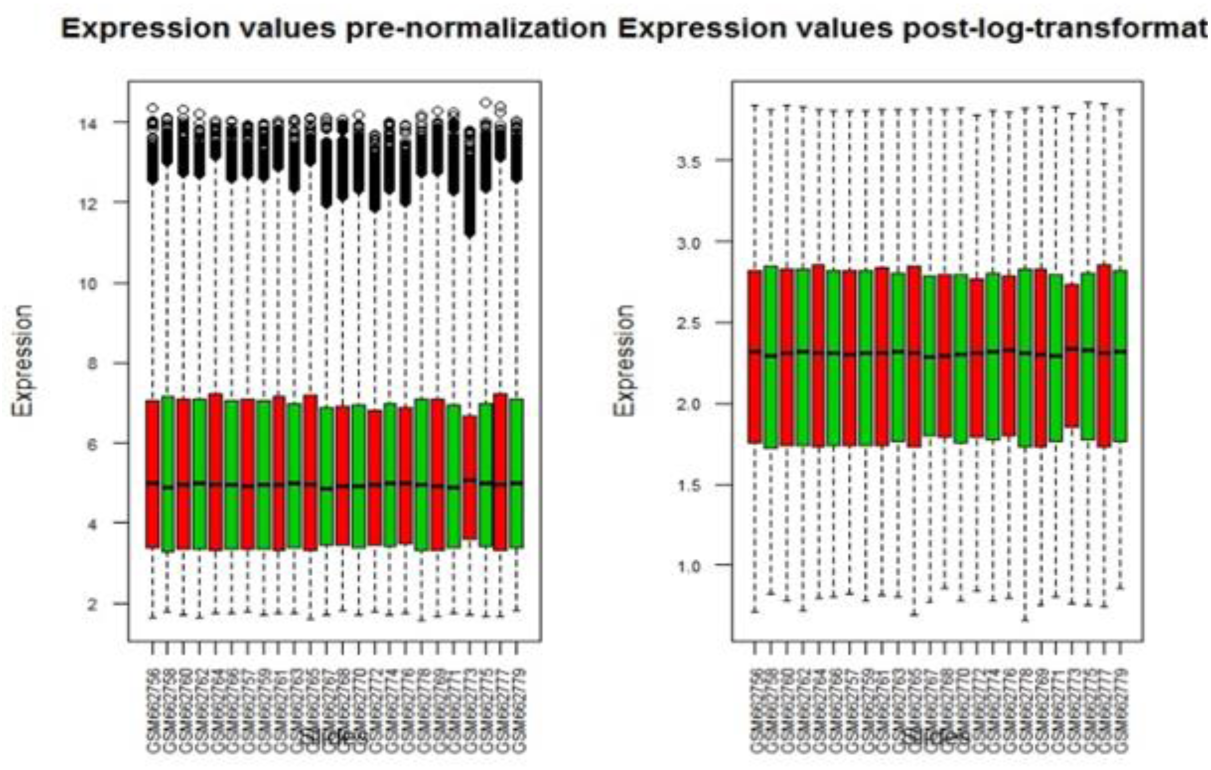
Box plots depicting sample expressions pre and post-normalization. Log 2 transformation is applied for the same and the data is rendered more balanced ahead of analysis. The cancer and non-cancer bars are rep-resented by red and green color codes.

### 2.3 Differentially Expressed Genes

With respect to the assumed significance level (α) to be 0.01 and hence a stern confidence interval of 99%, we aim to copiously optimize the gene(s) search by postulating as follows:

> *(Null hypothesis)H*_0_*: Genes are not differentially expressed (equal means)*
>
> *(Alternate hypothesis)H*_1_*: Genes are differentially expressed (unequal means)*

The listing of differentially expressed genes will implicitly catalogue up- and downregulated genes too. To prudently list them out, a within genes correlation does the job. The negative numbers represent down-regulated genes and positives up-regulated ones. (Danielsson et al. 2013) report that maximum of genes en route malignancy, are down regulated. This is not for reference, but only to mark. There is also to note that since breast cancer and prostate cancer find unique origins pertaining sex discrepancy, it’s only logical to work with bifurcated dataset. We contemplate breast cancer and prostate cancer entries with distinct exegesis and later combine and compare the results owing to significance to biological interpretation.

The exploration renders 356 probes being differentially expressed in breast stroma and 221 in prostate stroma (with p-value < 0.01) amongst which *ADH1B*, *COL10A1* are most distinctly expressed in breast strata, while *BMP5*, *SFRP4* are notable enough in prostate cluster…

**Figure 3.**
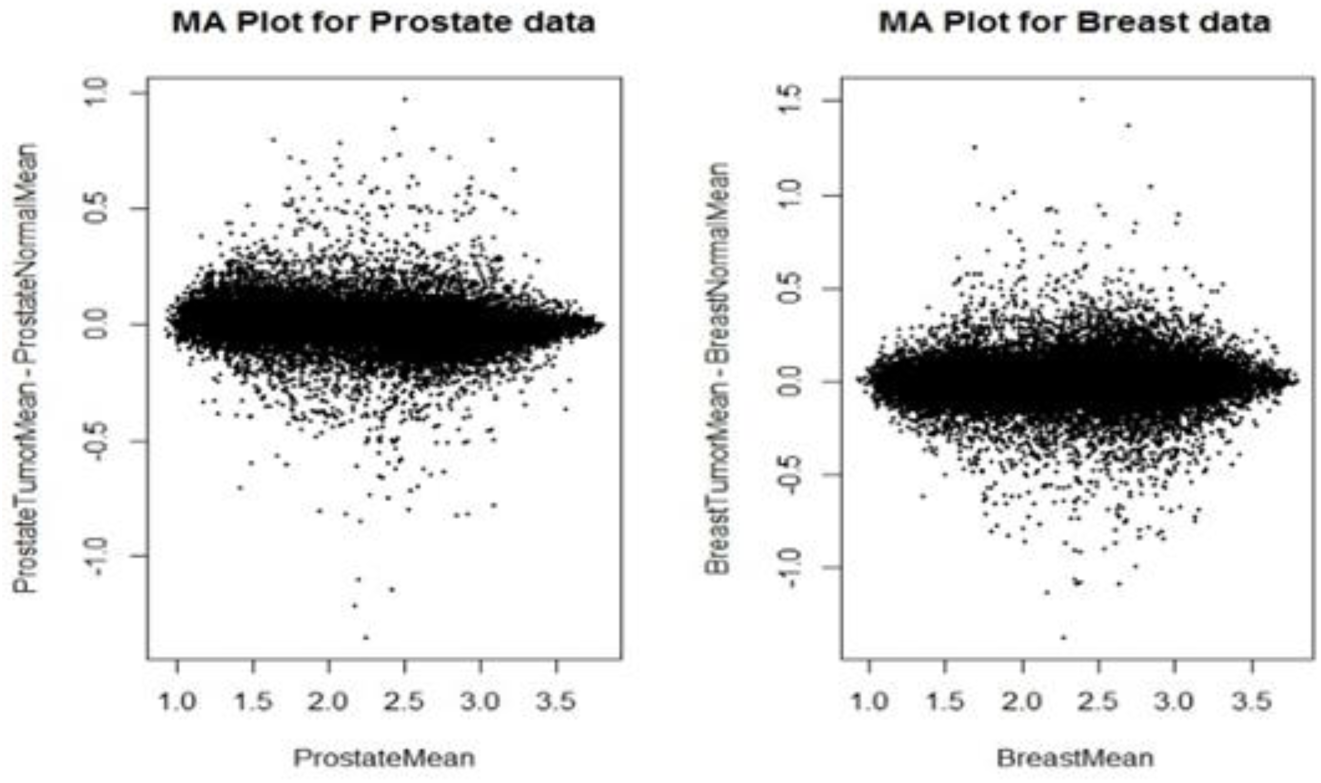
Respective MA plots of prostate and breast subsets. Clearly the floating specks demarcate the differentially expressed transcripts.

**Figure 4.**
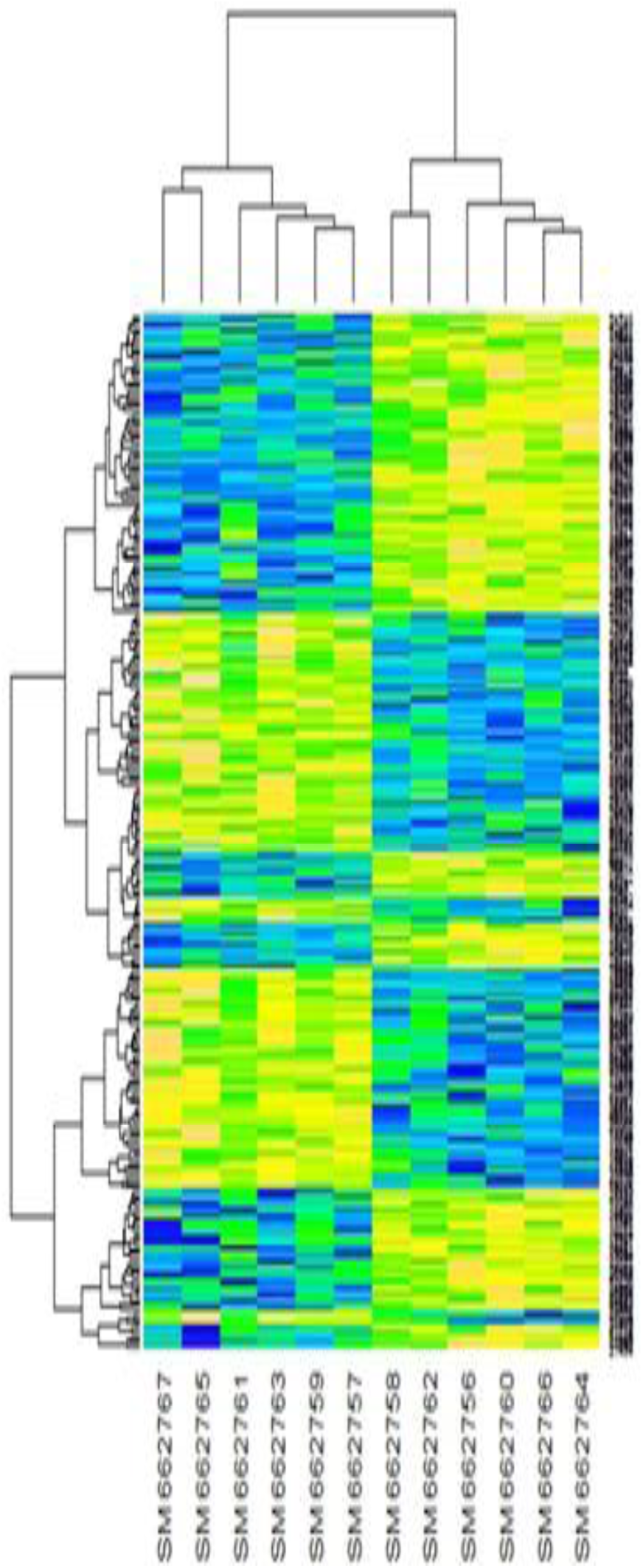
Heat map for differentially expressed breast genes. 356 probes with p-value < 0.01 were unraveled as being significant.

Desmoplastic Retort to Prostate and Breast Carcinomas’ Metastasis.

**Figure 5.**
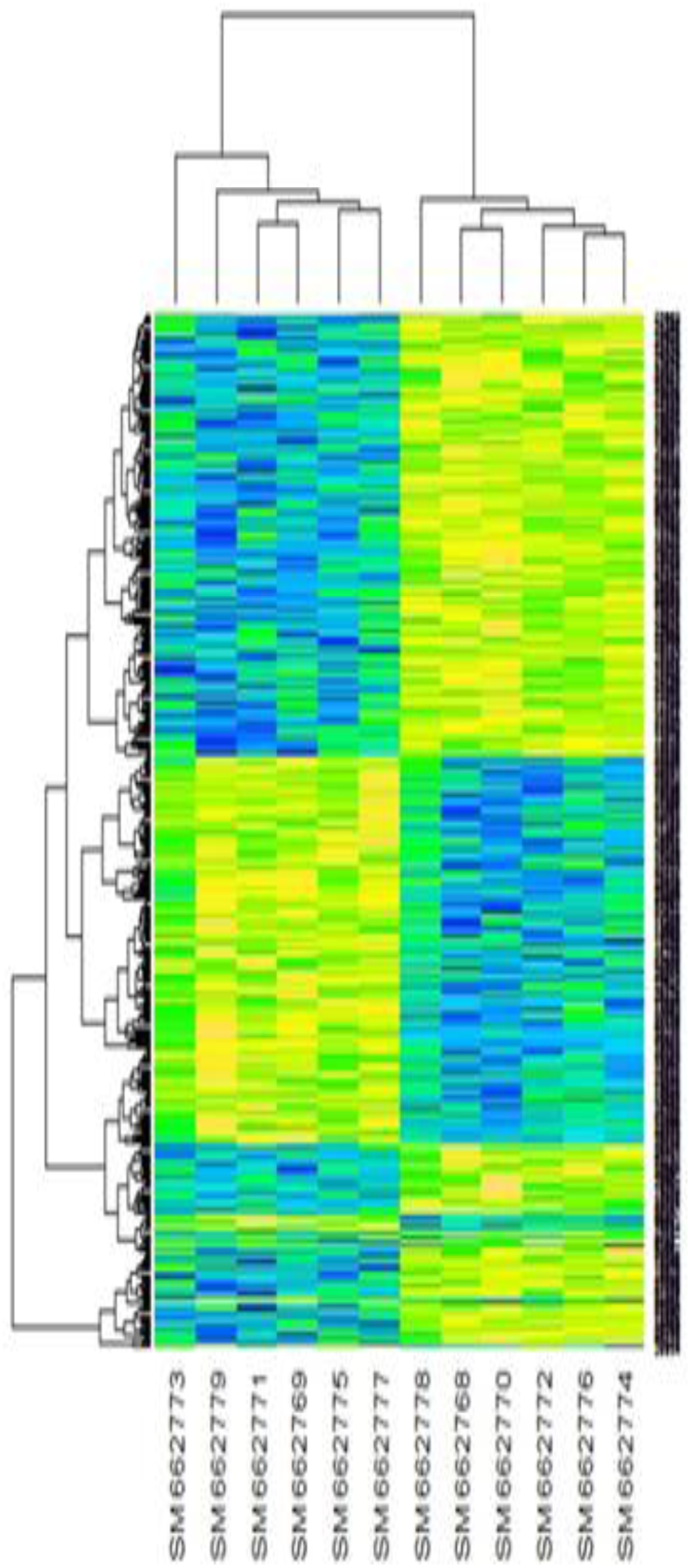
Heat map for differentially expressed prostate genes. Here, 221 probes with p-value < 0.01 were deemed crucial.

We also acknowledge that packages like *siggenes* (Schwender, 2012), *samr* are available that incorporate Significance Analysis of Microarrays (SAM) (Tusher, Tibshirani and Chu, 2001) working procedure, but in view of keeping the analysis more abstract and interactive, there is a minimum use of readymade library functions.

### 2.4 Gene Set Enrichment Analysis

**Figure 6.**
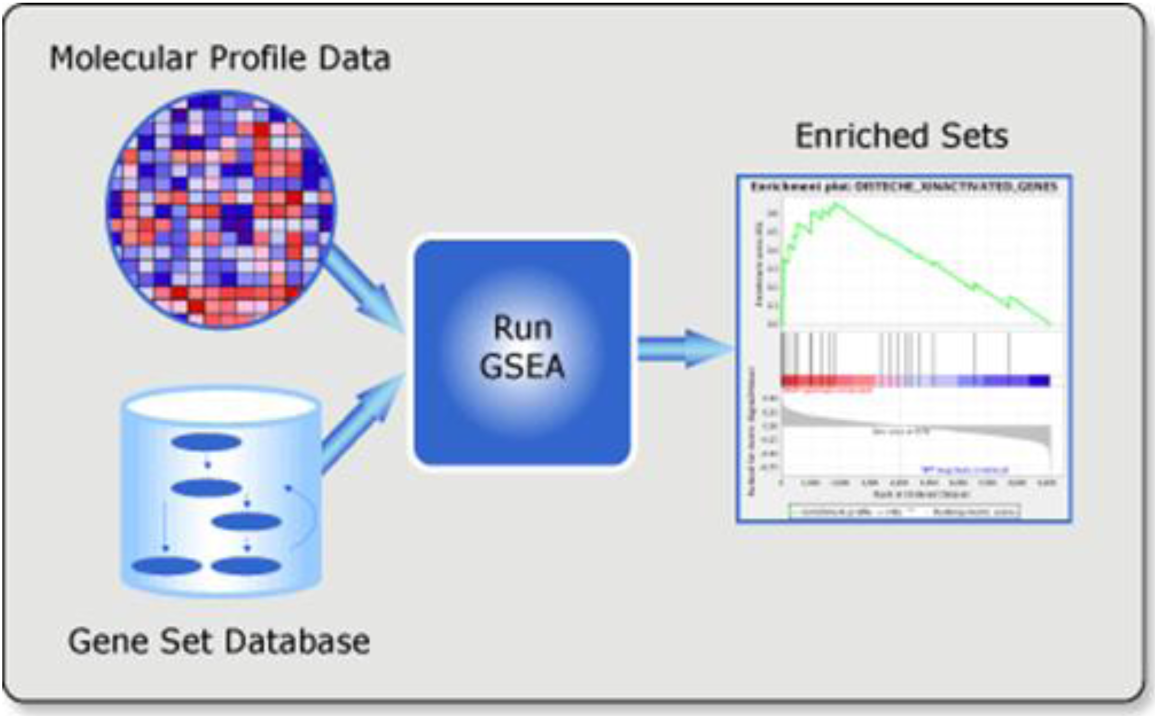
Illustration of GSEA framework (Subramanian, Tamayo, et al., 2005).

Gene Set Enrichment Analysis (GSEA) is a scheme to map statistically relevant genes to pre-known biological profiles, eg. phenotype, to discern their life relevance. The molecular signatures are updated as the curation cascades. There exist a consortium of metadata libraries for cataloguing genes and gene products’ information. To standardize the practice of annotation in genomics, this bioinformatics initiative is absolutely imperative as we’re riding the snowball of discoveries in GWAS. (In R language, GWAS is facilitated by Fischer’s exact test.)

This method is applied to the resultant set of differentially expressed genes and only those with a reference in MSigDB (Subramanian, Tamayo, et al., 2005) (with a valid Unigene_ID, Entrez_ID, and GO_TERM) are selected for further analysis. To accomplish the same, MeV and Cytoscape (in parts) are used. The GO listing wasn’t available for 10 probes in prostate data and 11 in breast data, which led to their discard. At this stage, our dataset has 345 and 211 rows in breast and prostate data, respectively.

### 2.5 Functionally Coherent Genes

An important aspect that escorts investigation of the differentially expressed genes is the strength of associativity between their tumor and normal roles. This can be explored using correlation technique of statistical descent. Commonly known Pearson’s product-moment (or simply, correlation) coefficient helps establish connect between two linearly distributed variables. In simplified terms, Spearman’s coefficient is a non-parametric version of Pearson’s coefficient with ranked data (Hauke & Tomasz 2011). Since, our dataset selection is so, we would prefer using Pearson’s correlation measure as opposed to Spearman’s or Kendall’s which is equally effective (or more) for the *qualitative* data. Kendall’s τ is based on concordance and discordance. The question is to establish similarity between two distinct genes, technically two expression vectors (Saeed et al. 2003). An expression vector spans expression values vide all featured experimental conditions. Albeit microarrays are not known to cater isoform expression detection as they are not absolute exhibitors of gene expression and rather give a relative value (log ratios of hybridization intensities).

The Pearson’s correlation coefficient (r) for class labels X and Y is mathematically represented as follows:

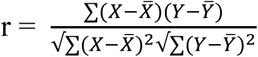

The resultant genes in breast cancer and prostate cancer were tested for correlation amongst themselves. With each gene confronting every other gene, a 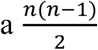 comparisons are expected in a pairwise matrix format. Since distance measure is hinged around mean values, a mix of positive and negative integers is likely. It is to note here that notwithstanding the amplitude of correlation, there is a significance of signs in the correlation matrix. A negative (-) number indicates that a gene is inhibiting another gene, while a positive (+) marks that the two are expressing collaterally. To safeguard our conviction to the fullest, the gene list was filtered with a dual-parameterized statistic. We sifted the genes with low p-value significance and high correlation measure. The tables catalogue genes from cancer-duos, with correlation > 0.95 and p-value < 10^-7^.

### 2.6 Gene Regulatory Networks (GRN)

Transcriptional activity can be precisely monitored with GRNs (Chai et al. 2014). A visualization of putative pathways and the absolute values that are symbolic of the degree of strength between two components can bring out some very useful linkage information. After the elucidation of differentially expressed genes, the inkling is to draw a correlation measure amongst them to infer a relational matrix with values {-1, 0, 1} with interpretations anticorrelated, no dependence, and correlated, respectively. This notion is certainly not delimited to “naivety of adjacency”. The informal theme is the distance measure, but logically it may falsify the overall outcome due to inherent biases with the chip construction. Therefore, the notion of correlated transcripts is revered more viable. A gene regulatory network is a visualization of a set/part(s) of genes that result in myriad (all) of cell processes, including metabolism, cell signaling and transduction, cell growth control etc., which is vital to understand the dynamics of molecular biology (Karlebach & Shamir 2008). But, the mechanistic inference of the architecture is subject to experimental biology, a wet lab gig (Davidson & Levin 2005). Nevertheless, the disposability of GRNs can’t be disparaged as they provide a blueprint of the underlying system and tellingly optimize our erudition.

Correlation establishes the linear propensity in-between variables. For pursuit of the same, we deliberate a Bayesian approach. In continuum to our expedition with R, the packages, BNArray (Chen et al. 2006), NATbox (Chavan et al. 2009) deploy probabilistic slant to decipher gene interactions, where NATbox shows competitive proficiency (Chavan et al. 2009). In this treatise, however, we’ve considered Cytoscape as a pliant tool for visualization the transcriptional network in corroboration with the *GeneMania* plugin. The output network matrix of genes was exported to Cytoscape for visualization and analysis.

Post validation of the transcriptional networks, gene ***CCDC11***, which has traditionally been revered for human laterality disorder (Perles et al. 2012; Narasimhan et al. 2015), has been elicited to show strong propensity in both (breast and prostate) cancer profiles. The **Coiled-Coil Domain Containing 11** or CCDC11 is a protein coding gene which is closely associated with epidermis in amphibians and skin fibroblasts from Homo sapiens (Narasimhan et al. 2015). Re-annotated as **Cilia and Flagella Associated Protein 53 (CFAP53)**, the mutation in CCDC11 exhibits perturbed left-right asymmetry (Silva et al. 2016). As showcased in the particular analysis, it has thorough connectedness at the order of 11 (aggregate) to other genes, which may signify functional co-regulation. From the understanding, it is proposed that this hub gene could be responsible to stage the process of stromal response and coordinate in the transcriptional activities of the same.

***WDR88*** gene on chromosome 19, revered WD repeat domain 88, is a protein-coding gene and a branded marker for the onset of prostate cancer (Chinnaiyan et al. 2013). In a topical finding, the gene has also been shown to have links with schizophrenia (Richards et al. 2016). 167 organisms have orthologs with human gene WDR88 that is conserved in chimpanzee, Rhesus monkey, dog, cow, mouse, rat, chicken, and frog.

**Figure 7.**
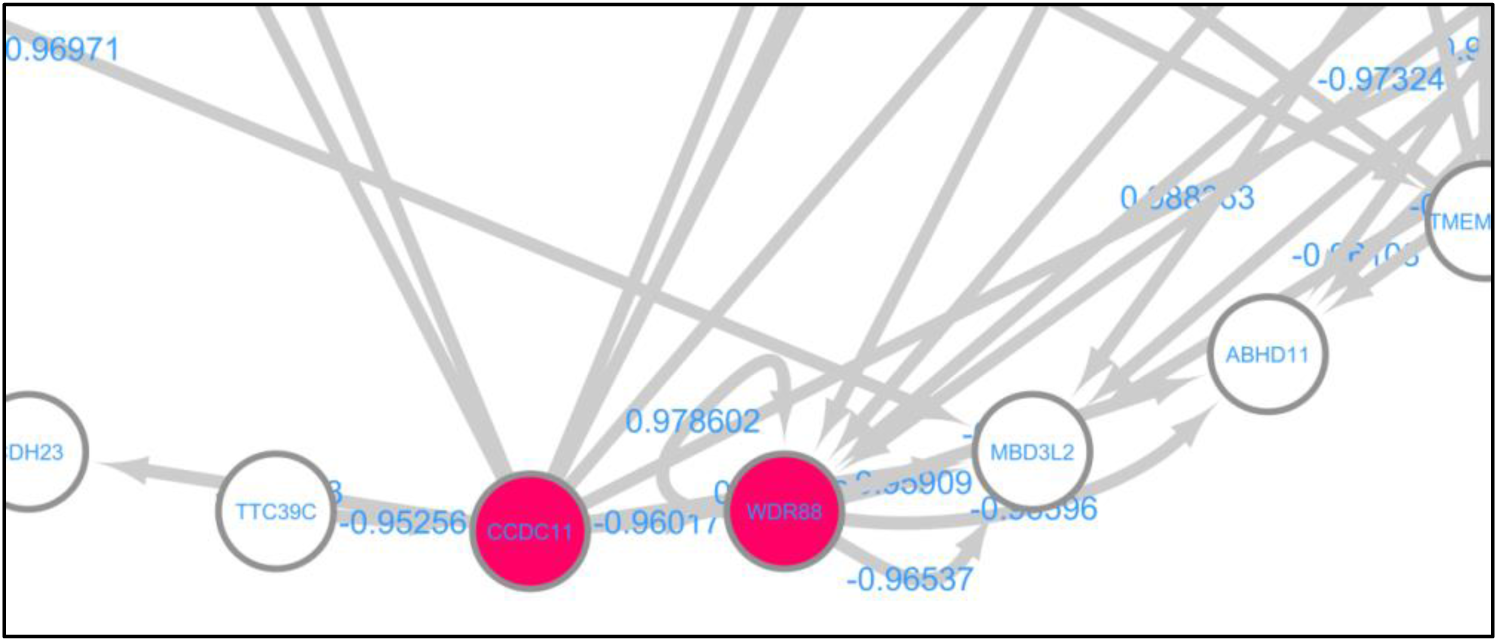
The study fortifies the eminent role of CCDC11 and WDR88 genes that are fundamental test genes for cancer diagnosis. The figure portrays a window from the prostate cancer GRN. As elicited, CCDC11 is orchestrating other genes, while WDR88 is a coveted.

Another gene ***ARPP21***, located in chromosome 9, has been exceptionally highlighted in the breast and prostate cancer profiles. It has been duly captured to be frequently deregulated as is miR-128 (Pellagatti et al. 2010; Li et al. 2013). According to NCBI RefSeq (June 2012), this gene encodes a cAMP-regulated phosphoprotein. The encoded protein is enriched in the caudate nucleus and cerebellar cortex. A similar protein in mouse may be involved in regulating the effects of dopamine in the basal ganglia. Alternate splicing results in multiple transcript variants. It is thence fathomed that these hub genes could be responsible to stage the process of stromal response and coordinate in the transcriptional activities of the same.

**Figure 8.**
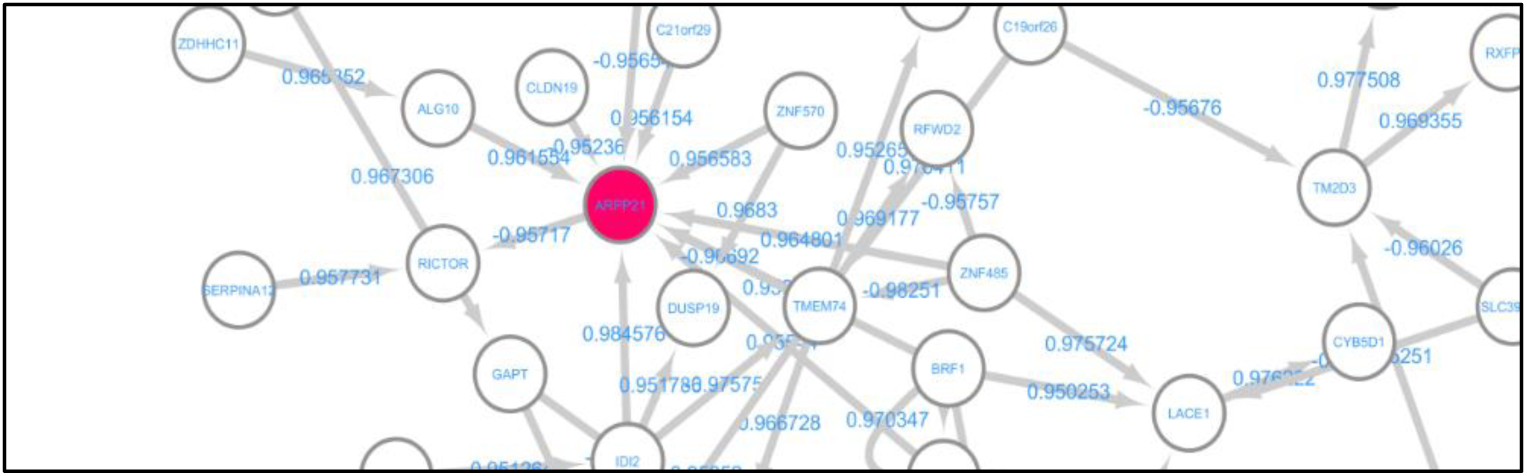
An excerpt from the Breast cancer GRN analysis shows profound coverage of ARPP21 gene with high propensity.

From gene ontology, ARPP21 is also attributed to the response to stimulus, triggering cellular response to heat; at any temperature higher than the optimal stimulus of that organism.

A deliberation to the current study also entails the exceptional, yet formidable idea of cross-linkages of breast and prostate cancers. Although the rudiments of breast and prostate are oriented towards females and males, respectively, nonetheless, an exceptional yet indelible facet of female prostate and male breast profiles has been dimly studied. According to the American Cancer Society, breast cancer is aggregate 100 times less common in males than females; that is to calibrate the lifetime risk of a male getting breast cancer is 1 in 1000. Contrastingly, the skene/ periurethral gland carcinoma (female prostate cancer, in generic terms) is also found to contribute less than 0.0003 percent towards all genital cancers in women (Dodson et al. 1994). The numbers aren’t intellectually stimulating, albeit we choose to delve a little deeper.

### 2.7 Female Prostate and Male Breast Carcinomas

Female prostate, i.e. Skene gland, named after Alexander Johnston Chalmers Skene, who was a British gynecologist from Scotland, is a homologue for the male prostate organ and its adenocarcinoma is a scarce occurrence. Elevated Prostate Specific Antigen (PSA) and PSA Phosphatase (PSAP) are potent markers for detecting prostate cancer in general (female as well as male prostate specimens). Owing to the rarity, the female prostate cancer isn’t thoroughly researched too. With the limited physiological understanding, an older case study presented a female subject with advanced form of the disease. It was treated with convention-al surgery (Ueda et al. 2012), to eventually weed out all the spread. Other techniques including radiotherapy (Korytko et al. 2012) have been sublimely effective as well.

The female prostate is acknowledged as a functional part of the urinary and reproductive systems in female humans (Zaviacic et al. 2000). It is located on the anterior wall of the vagina, around the lower end of urethra, on each side. A chance that skene gland could be a secondary cancer site is also plausible. Estrogen (Estradiol, Estriol, and Estrone) and Progesterone are the two key enzymes/ hormones that regulate the female breast development, menstrual cycle, and sexual function. They are also luminaries in the prostate region in the female gerbils. Estrogen is present in both male humans and female humans, and can be measured for analyzing cancer of the reproductive system subunits, viz. ovaries, testicles, etc. The cancer of the Skene gland is also more recognized in older females, showing tangible lesions (Custodio et al. 2010). Additionally, it has also been extensively deliberated that a family history of breast cancer and prostate cancer engenders augmented jeopardy to a postmenopausal woman gestating breast cancer (Robinson et al. 2015), (Beebe-Dimmer et al. 2015). The abscesses in the gerbils are also shown to be driven by progesterone. A case history of multiple pregnancies and ageing could be culpable for the female prostate disorder (Oliveira et al. 2011).

Owing to the limited case studies of Skene gland cancer, the symptoms of the disorder aren’t well acknowledged and etiology is apparently impervious. As general indications, bleeding in the urethra, that could also accompanied by sporadic pain, are primarily contingent to symptomatic treatments. If the following conditions hold, a quick visit to the physician is often advisable.

- Arduous, frequent, and often difficult urination
- Bleeding from the urethra
- Painful sexual intercourse and pubic area
- Erratic menstrual cycle

The causes for Skene gland cancer are diversely plethoric. They can include infection as prostatitis, some sexually transmitted infections (STIs) as gonorrhea; Polycystic Ovarian Syndrome (PCOS) that renders imbalance and frequently abundance of reproductive hormones, cysts, and adenofibroma.

**Figure 9.**
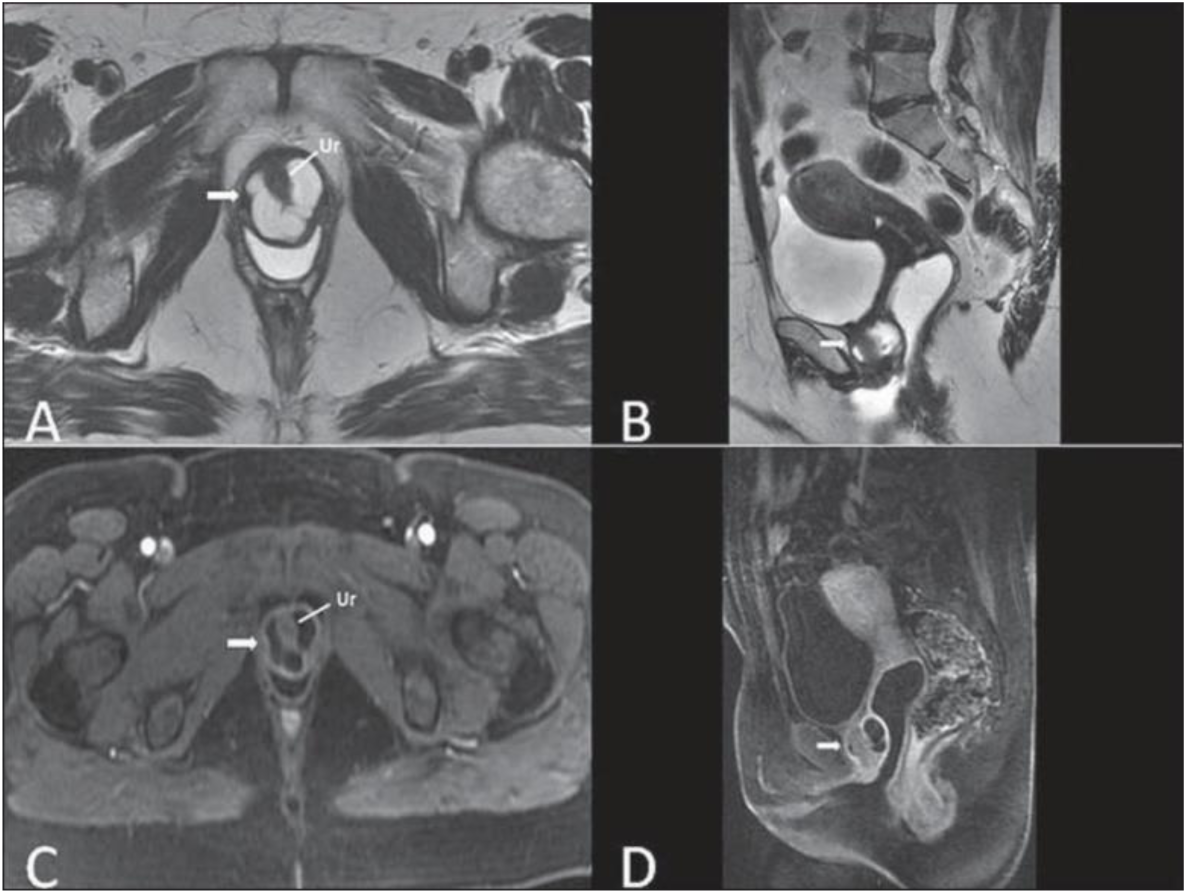
MRI scans of the cysts of the Skene glands. Multiplanar MRI T2-weighted (A,B) and contrastenhanced T1-weighted (C,D) sequences identifying distal periurethral cysts (Ur) (arrows) locat-ed between the urethra and the vagina.

Another malady, although uncommon but not to be belittled as the rate of occurrence increases every year, is the Male Breast Cancer (MBC). Mainly, females are more vulnerable to breast cancer, having stocky breast tissue; however, males have pertinent breast tissue as well. Scientifically, mutated copies of *BRCA1* and *BRCA2* genes incubated by male humans are proverbial causes for MBC. Tamoxifen and anti-hormonal drugs are FDA-approved chemotherapeutics to treat breast cancer in both male and females. Requisite surgery (mastectomy/ lumpectomy) followed by radiation therapy is standardly warranted, although individual therapies could include more aggressive treatment options. The ideology of a male human incubating breast cancer is largely pondered with ignorance and aversion; this conviction, in most cases, delays the screening of the disease. Peculiar symptoms of MBC entail ruptured (and often painful) nipples, puckering and dimpled masses of the breast, decolorized jaggy surfaces, etc. MBC is usually detected as a hard lump underneath the nipple and areola. The histopathological derivatives in MBC and female breast cancer are homogenous.

**Figure 10.**
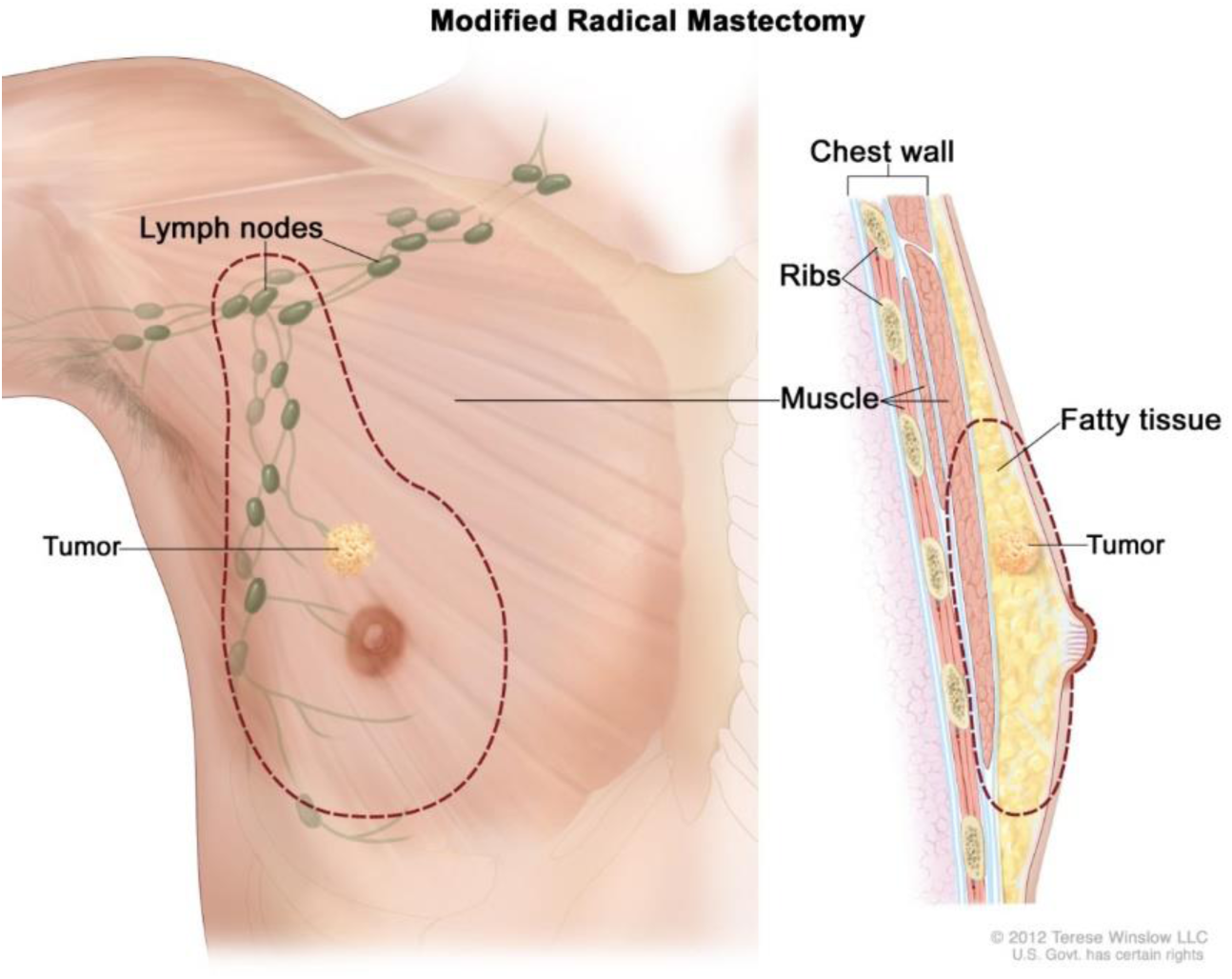
An illustration detailing radical mastectomy. Credit: http://www.cancer.gov.

**Figure 11.**
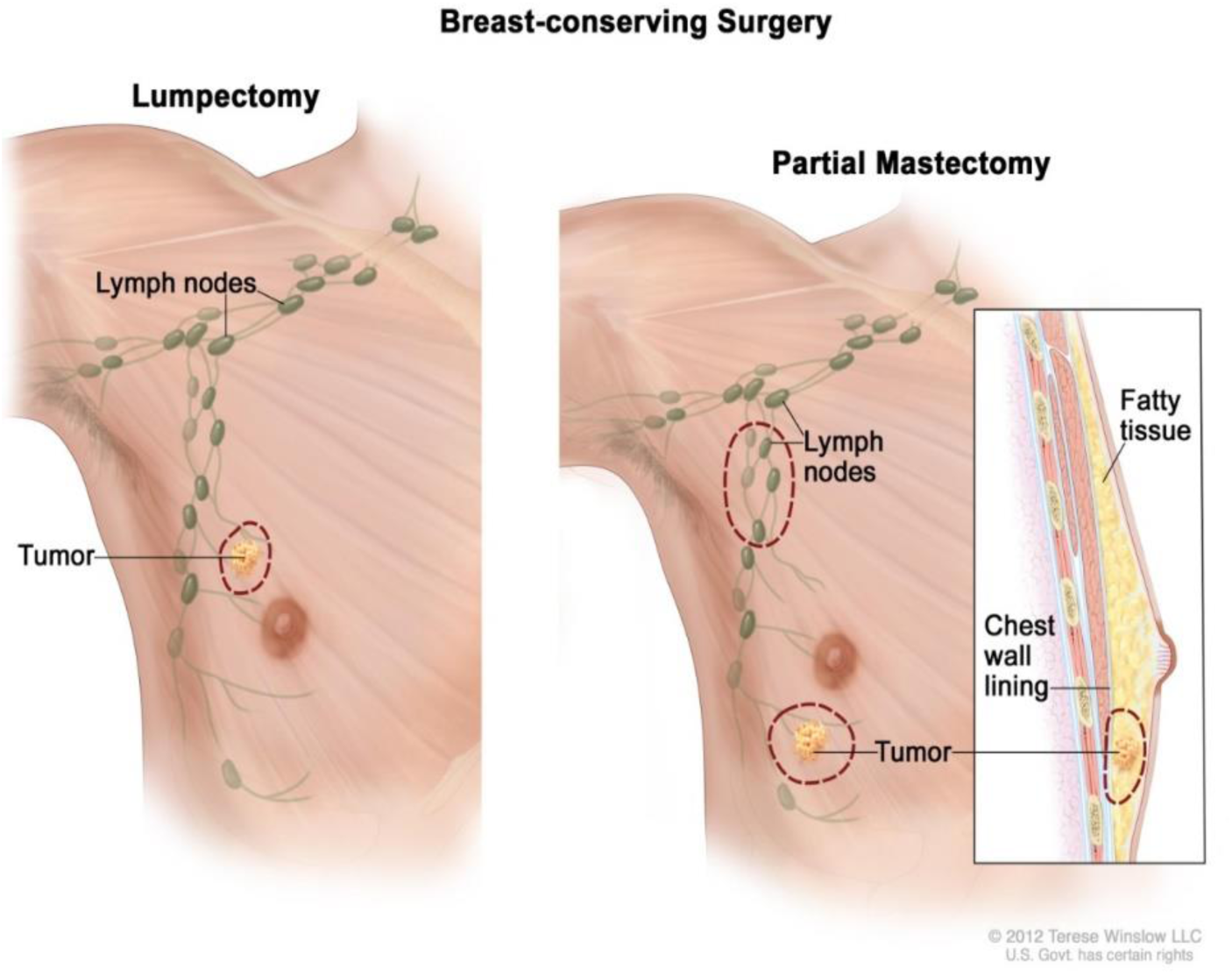
An illustration detailing breast-conserving surgery. Credit: http://www.cancer.gov.

Gynecomastia is also a disorder in men of benign nature, where the breast tissue becomes enlarged due to hormonal misbalance (oestrogen to testosterone ratio), especially during puberty. Although a natural phenomenon, it is usually conceived with humiliation and anxiety; however paltry cases have been reported to establish that gynecomastia and MBC are concomitant. In conjunction, pseudogynecomastia is a condition when adipose tissue (fat) causes gynecomastia.

Therefore, it can be argued that the denominations of origins, histopathology, causes, symptoms, and treatments are overlapped for male-breast and female-breast cancer; and likely so for male-prostate and female-prostate cancer. The contributing genes and pathways could be further explored for overlap in disease profile and therapy.

### 2.8 Stromal Response Threshold

The cancer metastasis presents an intriguing case of classical science theory- a medium required for propagation. The carcinogenesis triggers a parallel desmoplastic reaction that serves as carrier of malignancy (Whatcott et al. 2012). Technically, desmoplasia is the development of fibrous and connective tissues encompassing tumor cells. Cells like endothelium and fibroblasts stage all structural and functional profiles from carcinogenesis to metastasis (Kalluri & Zeisberg 2006) and vitally graded therapeutic targets. Chemo resistance is highly attributed to mutations in cancer cells as per the Darwinian doctrine of evolution, survival of the fittest (Fodale, Pierobon, Liotta, & Petricoin, 2011) (Pisco et al. 2013).

We import widely recognized *e1071* package library (Meyer et al. 2015) to employ Support Vector Machine (SVM) classification to distill the transcriptional threshold to desmoplastic response. Although there are four others catalogued in the R library that carry out the SVM implementation, viz. *kernlab*, *klaR*, *svmpath*, and *shogun*. Technically, a decision boundary equation is sought here. Our aim, from epidemiological context, is to aid medicinal normalization of transcriptional impressions so as to contain tumor invasion trans-organismal cultures. The one-versus-one favor of classification is evident for 6 class pairs (normal-tumor duo). Multiple kernel types were considered and cost functions analyzed before arriving at the *tune()* that cross-validates a range of SVM models outputs the optima.

The equation of the hyperplane separating the negative and positive examples is given by:

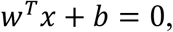

where *w* is weight vector, *x* is input vector, and *b* is bias. The decision boundary can also be deemed as a linear combination of support vectors. As calculated, 8 and 9 support vectors were rendered from prostate and breast data respectively. The bias vectors from *<svm model>$rho* are 1.429841 and 4.139861, from prostate and breast data correspondingly. Further information can be found, as code output, from the supplementary documents.

**Figure 12.**
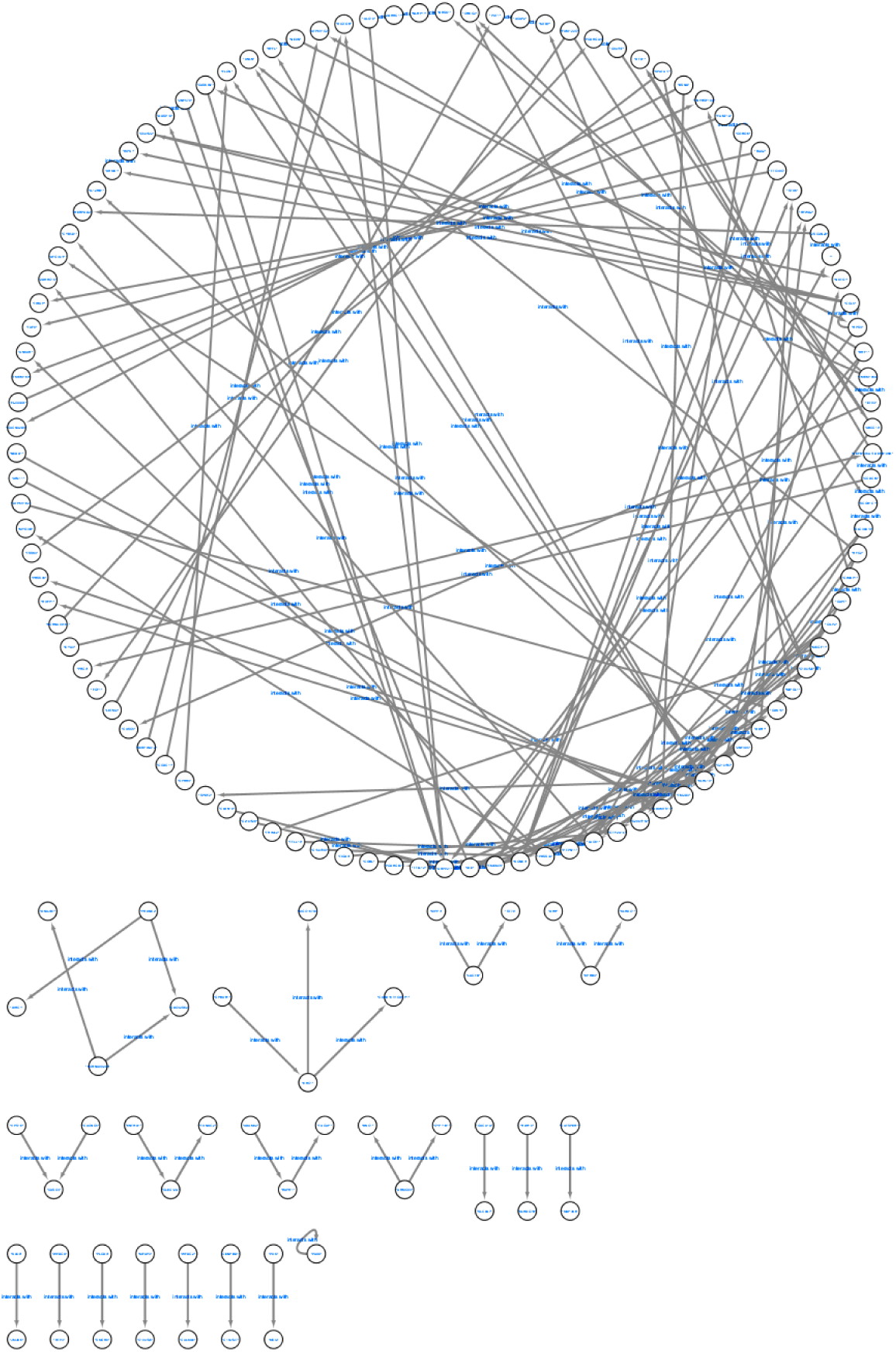
Breast Genes Regulatory Network.

**Figure 13.**
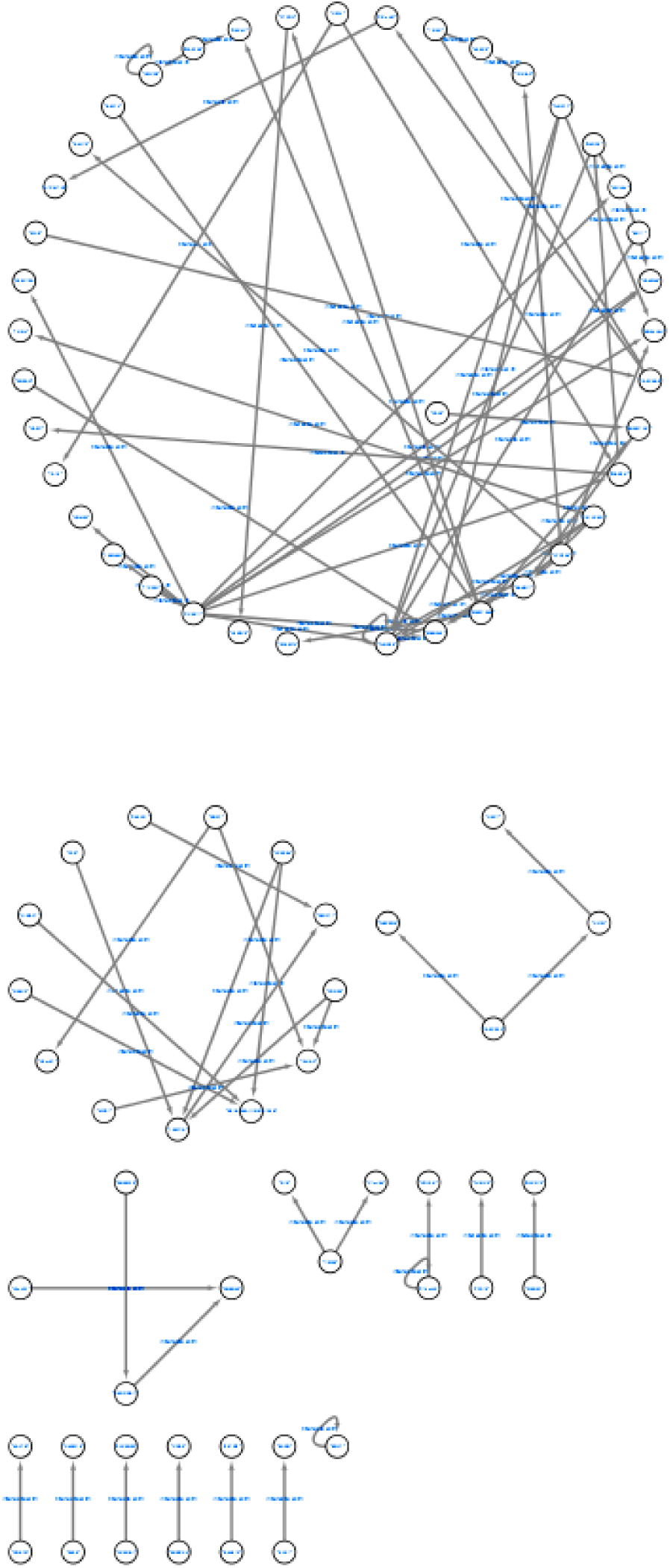
Prostate Cancer Regulatory Network.

### 2.9 Conclusion and Future Work

This text has been premeditated to render the most interactive portrayal of working with gene expression data analysis. As a part of the original work, the authors have carried out survival analysis too. The treatise however concentrates on the improved biomarker(s) dissemination.

As an imminent applicability, the study can aid fostering of pertinent therapeutics to deride proliferation of cancer metastasis from one tissue to another by monitoring the expression threshold and keeping it checked.

The procedure highlights an illustration of the packages available in the R language and Bioconductor that duly facilitate the exploratory analysis of the genomic data. While doing so, certain cohorts of genes were found relevant and were statistically narrowed to seed further analysis. This aids reducing the search space for biomarkers (broadly explains the doc-trine of bioinformatics) and the pipeline of wet laboratory testing and validation, proceeds. If the genes *CDCC11*, *WDR88* and *ARPP21* have any causal implications in the stromal response to the cancer metastasis, can and will only be substantiated through valid laboratory studies. Researches alike add to the annotations of the known gene functionality. An array of such explorations is warranted and is indeed happening. This trend over a period of time is believed to pave way for a precision medicine schedule, when drug compounds’ applications and the respective gene functions are almost perfectly matched.

## 3. Material and Methods

### 3.1 Dataset Selection

The gene expression dataset chosen from the study is derived out of a study based on stromal cells and invasive breast and prostate cancer development (Planche et al. 2011)

**Table 1.** Dataset Profile.

| S. No. | Parameter | Value |
| --- | --- | --- |
| 1 | Sample Count | 24 |
| 2 | Value Type | Transformed<br>Count |
| 3 | Channel Count | 1 |
| 4 | Platform Organism | Homo sapiens |
| 5 | Platform | *In situ* |
|  | Technology | *oligonucleotide* |
| 6 | Sample Type | RNA |
| 7 | Feature Count | 54675 |
| 8 | Dataset Platform | GPL570 |
| 9 | Dataset<br>identification | GDS4114 |
| 10 | Series | GSE26910 |

**Table 2.**
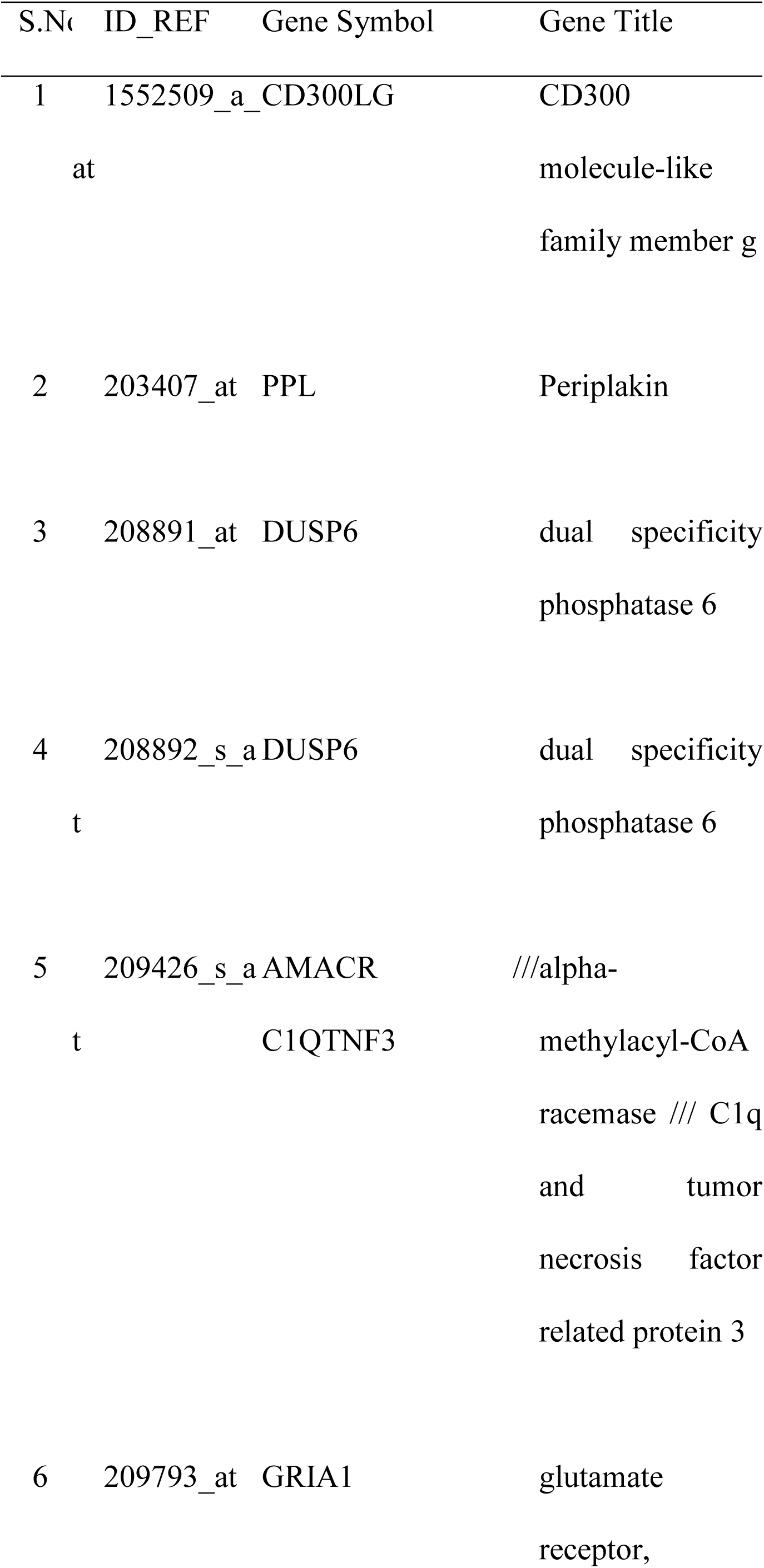
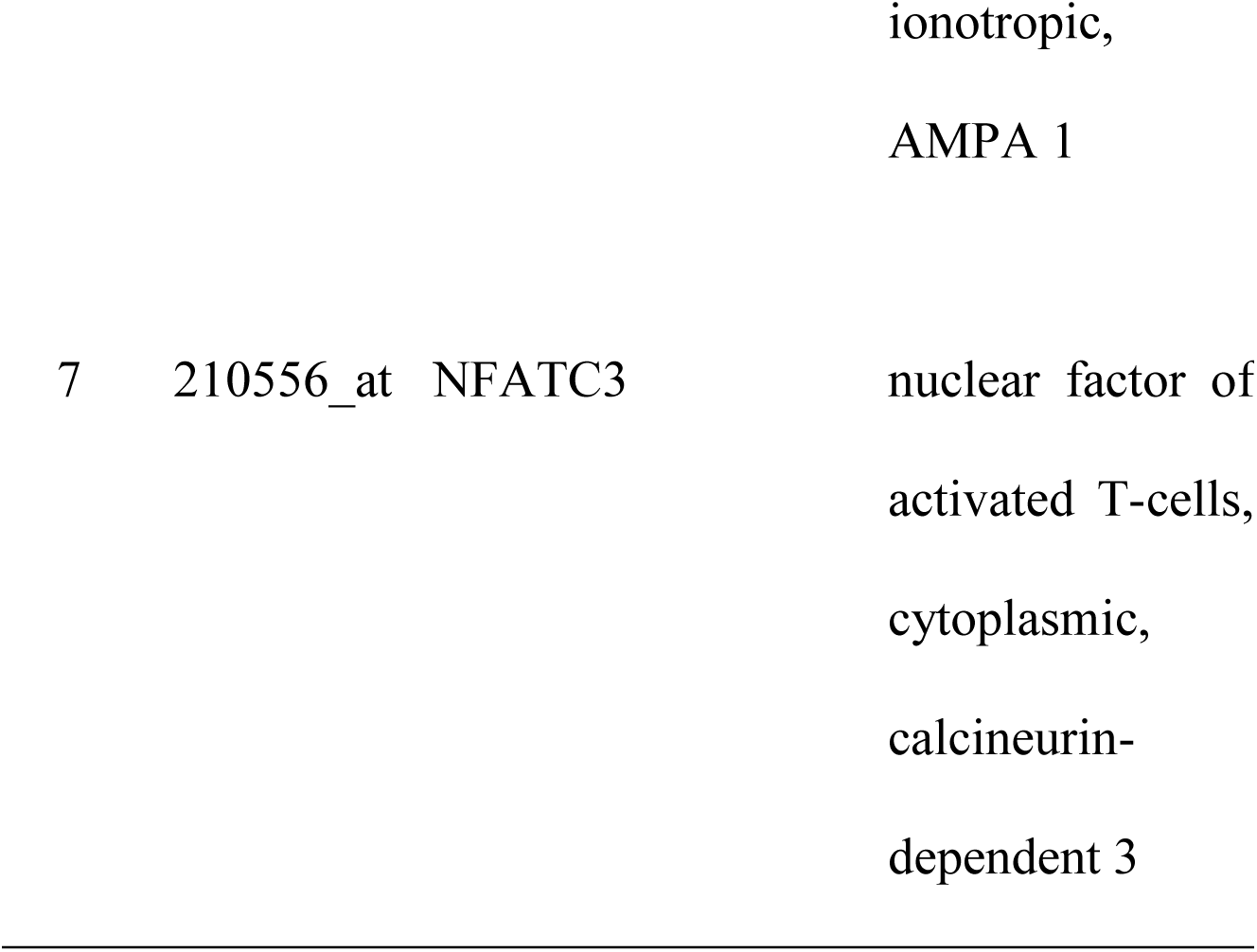
Intersecting transcripts in breast and prostate data.

**Table 3.** Breast cancer network visualization ready tabulation.

| Source/From | Target/To | Correlation | P-value |
| --- | --- | --- | --- |
| Gene | Gene |  |  |
| PAX8 | PAX8 | -0.96158 | 6.18E-07 |
| DDR1 | AFG3L1P | -0.96217 | 5.72E-07 |
| ZDHHC11 | ALG10 | 0.965852 | 3.45E-07 |
| C15orf40 | PRSS33 | 0.957133 | 1.06E-06 |
| TTC39C | TIRAP | -0.96351 | 4.79E-07 |
| PXK | MSI2 | -0.95376 | 1.54E-06 |
| CORO6 | FAM71A | -0.95738 | 1.03E-06 |
| GIMAP1 | GAPT | 0.967306 | 2.78E-07 |
| SPATA17 | TSSK3 | 0.974818 | 7.64E-08 |
| ENTHD1 | CLEC12A | 0.955864 | 1.22E-06 |
| CENPBD1 | C15orf27 | 0.970408 | 1.70E-07 |
| WFDC2 | CALML6 | 0.962181 | 5.72E-07 |
| EYA3 | DEFB106A /// | 0.953635 | 1.56E-06 |
|  | DEFB106B |  |  |
| CCDC65 | DEFB106A /// | 0.970579 | 1.65E-07 |
|  | DEFB106B |  |  |
| MFAP3 | C10orf25 | 0.958966 | 8.55E-07 |
| TMEM106A | ETV3 | 0.955541 | 1.27E-06 |
| KLHL10 | KLHL10 | 0.969253 | 2.06E-07 |
| RFC2 | TM2D3 | -0.95251 | 1.76E-06 |
| SLC39A13 | TM2D3 | -0.96026 | 7.30E-07 |
| C19orf26 | TM2D3 | -0.95676 | 1.11E-06 |
| PRSS33 | ANKAR | 0.956311 | 1.16E-06 |
| SLC39A13 | SCGB1C1 | 0.952224 | 1.81E-06 |
| CCDC65 | SCGB1C1 | -0.96076 | 6.86E-07 |
| FAM122C | SCGB1C1 | 0.972535 | 1.18E-07 |
| ADAM32 | RAPH1 | -0.96305 | 5.10E-07 |
| PLCD3 | SMCR8 | -0.96539 | 3.69E-07 |
| ARMCX4 | BNC1 | -0.95534 | 1.30E-06 |
| LACTB | MPP4 | 0.958924 | 8.59E-07 |
| DDR1 | LACE1 | 0.952876 | 1.69E-06 |
| SLC39A13 | LACE1 | -0.95083 | 2.08E-06 |
| BRF1 | LACE1 | 0.950253 | 2.21E-06 |
| ZNF485 | LACE1 | 0.975724 | 6.37E-08 |
| TTLL12 | IDI2 | 0.951264 | 1.99E-06 |
| ARMCX4 | CYP11B1 | 0.972441 | 1.20E-07 |
| RAX2 | TAF8 | -0.95835 | 9.20E-07 |
| TMEM106A | BRSK1 | -0.95993 | 7.61E-07 |
| TTC39C | KCNE4 | 0.965008 | 3.90E-07 |
| CILP2 | KCNE4 | -0.95681 | 1.10E-06 |
| RAX2 | KCNE4 | -0.95326 | 1.62E-06 |
| COBL | KCNE4 | 0.950202 | 2.22E-06 |
| CCL5 | HIPK1 | 0.952769 | 1.71E-06 |
| C19orf26 | MTBP | 0.966859 | 2.98E-07 |
| ACAP2 | MTBP | 0.95023 | 2.21E-06 |
| C19orf26 | TMEM74 | 0.970411 | 1.70E-07 |
| ZNF485 | TMEM74 | -0.98251 | 1.25E-08 |
| IDI2 | TMEM74 | -0.97575 | 6.34E-08 |
| IDI2 | C21orf67 | 0.950287 | 2.20E-06 |
| TMEM74 | C21orf67 | -0.95852 | 9.02E-07 |
| ZDHHC11 | NLRP11 | 0.950934 | 2.06E-06 |
| BRSK1 | NLRP11 | -0.9579 | 9.70E-07 |
| CCDC11 | ATP6V1C2 | -0.95934 | 8.17E-07 |
| BRF1 | ATP6V1C2 | -0.95839 | 9.16E-07 |
| RAPH1 | TAGAP | -0.95451 | 1.42E-06 |
| TM2D3 | PRSS36 | 0.977508 | 4.37E-08 |
| KCNE4 | ZDHHC15 | 0.966886 | 2.97E-07 |
| ATP6V1E2 | CDK15 | 0.958811 | 8.71E-07 |
| PRSS33 | CDK15 | -0.96044 | 7.14E-07 |
| GAPT | CDK15 | 0.956134 | 1.19E-06 |
| IDI2 | CDK15 | 0.950309 | 2.19E-06 |
| ZSCAN20 |  | 0.956726 | 1.11E-06 |
| TMEM74 |  | 0.952653 | 1.73E-06 |
| WFDC9 | RTP3 | 0.950728 | 2.10E-06 |
| RFC2 | MIPOL1 | 0.951615 | 1.93E-06 |
| SPATA17 | MIPOL1 | 0.953921 | 1.51E-06 |
| MEGF11 | MIPOL1 | 0.954858 | 1.37E-06 |
| PRSS33 | MYO3B | -0.95153 | 1.94E-06 |
| MEGF11 | TRIML2 | 0.951737 | 1.90E-06 |
| C8orf47 | ABCC13 | 0.950995 | 2.05E-06 |
| C21orf67 | IL12RB1 | -0.97209 | 1.27E-07 |
| CILP2 | GTF2A1L | 0.956402 | 1.15E-06 |
| PRSS33 | GTF2A1L | 0.953581 | 1.57E-06 |
| MEGF11 | GTF2A1L | 0.952955 | 1.68E-06 |
| TIGD4 | GTF2A1L | -0.97121 | 1.48E-07 |
| GTF2A1L | GTF2A1L | 0.972378 | 1.21E-07 |
| ACAP2 | PXT1 | 0.951524 | 1.94E-06 |
| LETM2 | PXT1 | 0.950738 | 2.10E-06 |
| PRUNE2 | CDC42SE2 | -0.97319 | 1.04E-07 |
| NCRNA00204 | CDC42SE2 | 0.95647 | 1.14E-06 |
| ZNF485 | RFWD2 | -0.95757 | 1.01E-06 |
| TMEM74 | RFWD2 | 0.969177 | 2.08E-07 |
| DDR1 | STX6 | -0.96476 | 4.03E-07 |
| KCNE4 | STX6 | -0.95893 | 8.59E-07 |
| RFC2 | ANLN | -0.96962 | 1.94E-07 |
| KLHL10 | ANLN | 0.954726 | 1.39E-06 |
| FAM71A | TMEM163 | -0.95918 | 8.33E-07 |
| ZSCAN20 | HERPUD2 | -0.95071 | 2.11E-06 |
| KLK8 | JMJD6 | 0.953033 | 1.66E-06 |
| CATSPER1 | MAP3K6 | 0.983465 | 9.47E-09 |
| CILP2 | PTPN11 | 0.961084 | 6.58E-07 |
| PRSS33 | PTPN11 | 0.95872 | 8.81E-07 |
| TTLL10 | PTPN11 | 0.951342 | 1.98E-06 |
| MEGF11 | PTPN11 | 0.957441 | 1.02E-06 |
| MIPOL1 | PTPN11 | 0.953438 | 1.59E-06 |
| ABCC13 | PTPN11 | 0.975157 | 7.15E-08 |
| RDH10 | KLHDC7B | 0.960543 | 7.05E-07 |
| CCDC65 | PHC3 | 0.976601 | 5.31E-08 |
| GIMAP1 | TNFRSF10A | 0.976163 | 5.82E-08 |
| FLJ30901 | TNFRSF10A | 0.971343 | 1.45E-07 |
| KLHL10 | RFFL | 0.950583 | 2.14E-06 |
| NEXN | RFFL | 0.955226 | 1.31E-06 |
| NEXN | HPS4 | 0.97026 | 1.74E-07 |
| HPS4 | HPS4 | 0.970347 | 1.72E-07 |
| CCL5 | UHMK1 | 0.972703 | 1.14E-07 |
| HIPK1 | UHMK1 | 0.955461 | 1.28E-06 |
| CLEC12A | TXNDC2 | 0.96067 | 6.94E-07 |
| CCL5 | C5orf22 | 0.956595 | 1.13E-06 |
| IDI2 | PCDHGB7 | 0.950937 | 2.06E-06 |
| GPBAR1 | ERC1 | 0.958682 | 8.85E-07 |
| KLHL10 | FLCN | 0.950509 | 2.15E-06 |
| SPRR4 | FLCN | 0.970464 | 1.68E-07 |
| GUCA1A | SLC9A7 | 0.956361 | 1.16E-06 |
| PRUNE2 | DIRC1 | 0.967442 | 2.73E-07 |
| NCRNA00204 | DNAJB7 | 0.953422 | 1.60E-06 |
| ETV3 | CASC5 | 0.950507 | 2.15E-06 |
| DDR1 | C20orf152 | -0.95261 | 1.74E-06 |
| CORO6 | C20orf152 | 0.957301 | 1.04E-06 |
| GAPT | C20orf152 | -0.96149 | 6.25E-07 |
| IDI2 | C20orf152 | -0.96458 | 4.14E-07 |
| TMEM74 | C20orf152 | 0.966728 | 3.04E-07 |
| TMEM106A | CASKIN1 | 0.985543 | 4.85E-09 |
| BSND | CACNA2D4 | -0.95795 | 9.65E-07 |
| ERC1 | MGC16703 | 0.954553 | 1.41E-06 |
| ERC1 | CARD16 /// | 0.954911 | 1.36E-06 |
|  | CASP1 |  |  |
| IDI2 | DUSP19 | 0.951786 | 1.89E-06 |
| ZNF570 | DUSP19 | 0.9683 | 2.39E-07 |
| LACE1 | CYB5D1 | 0.976222 | 5.75E-08 |
| C19orf26 | CREG2 | 0.967519 | 2.70E-07 |
| ETV3 | CREG2 | 0.957063 | 1.07E-06 |
| TM2D3 | RXFP1 | 0.969355 | 2.02E-07 |
| KLHL10 | SPEF2 | -0.95268 | 1.73E-06 |
| DEFB106A /// | DTD1 | 0.951094 | 2.03E-06 |
| DEFB106B |  |  |  |
| ABCC13 | DTD1 | 0.957523 | 1.01E-06 |
| VPS18 | CASC4 | -0.96446 | 4.21E-07 |
| CACNG5 | CASC4 | 0.953388 | 1.60E-06 |
| FAM122C | FGF1 | 0.953203 | 1.63E-06 |
| ALG10 | ARPP21 | 0.961554 | 6.20E-07 |
| BRF1 | ARPP21 | 0.958093 | 9.49E-07 |
| ZNF485 | ARPP21 | 0.964801 | 4.01E-07 |
| IDI2 | ARPP21 | 0.984576 | 6.70E-09 |
| TMEM74 | ARPP21 | -0.96692 | 2.95E-07 |
| CLDN19 | ARPP21 | -0.95236 | 1.78E-06 |
| ZNF570 | ARPP21 | 0.956583 | 1.13E-06 |
| C21orf29 | ARPP21 | 0.956154 | 1.19E-06 |
| HPS4 | ARPP21 | 0.95544 | 1.28E-06 |
| CASKIN1 | ARPP21 | -0.95654 | 1.14E-06 |
| NEDD1 | ADAMTS17 | 0.965781 | 3.49E-07 |
| GIMAP1 | ADAMTS17 | -0.95222 | 1.81E-06 |
| BSND | ADAMTS17 | -0.96552 | 3.62E-07 |
| ARL11 | ADAMTS17 | 0.964328 | 4.28E-07 |
| ADAMTS17 | ADAMTS17 | 0.953315 | 1.61E-06 |
| EPHB3 | DHH | -0.95225 | 1.80E-06 |
| EPHB3 | KLHDC1 | -0.96504 | 3.88E-07 |
| SERPINA12 | RICTOR | 0.957731 | 9.90E-07 |
| ARPP21 | RICTOR | -0.95717 | 1.06E-06 |
| C8orf47 | PCDHGA4 | -0.96422 | 4.35E-07 |
| NCRNA00161 | PCDHGA4 | -0.96077 | 6.85E-07 |
| C21orf67 | NETO1 | -0.95729 | 1.04E-06 |
| C5orf22 | NETO1 | 0.9569 | 1.09E-06 |
| LACTB | ST7L | -0.95159 | 1.93E-06 |

**Table 4.** Prostate cancer network visualization ready tabulation.

| Source/From<br>Gene | Target/To<br>Gene | Correlation | P-value |
| --- | --- | --- | --- |
| BEST4 | TMEM106A | -0.96094 | 6.70E-07 |
| CYP2A6 | ALG10 | 0.959298 | 8.22E-07 |
| C15orf40 | C15orf40 | 0.982069 | 1.42E-08 |
| TTC39C | CCDC11 | -0.95256 | 1.75E-06 |
| CYP2A6 | TRIOBP | -0.95726 | 1.05E-06 |
| C15orf40 | CRYZL1 | -0.95591 | 1.22E-06 |
| TRIOBP | LEAP2 | 0.951457 | 1.96E-06 |
| TIRAP | LEAP2 | 0.970564 | 1.66E-07 |
| FAM122C | SCIN | -0.97165 | 1.38E-07 |
| TIRAP | FAM18B2 | -0.95748 | 1.02E-06 |
| MSI2 | FAM18B2 | -0.97047 | 1.68E-07 |
| PRR22 | FAM71A | 0.957059 | 1.07E-06 |
| PXK | FAM71A | 0.957061 | 1.07E-06 |
| CCDC65 | FAM71A | -0.95069 | 2.11E-06 |
| SCIN | GAPT | 0.960974 | 6.68E-07 |
| DDR1 | C8orf47 | 0.953643 | 1.56E-06 |
| TIMD4 | C1orf65 | -0.95672 | 1.11E-06 |
| CCDC11 | CLEC12A | -0.95066 | 2.12E-06 |
| CCDC11 | CALML6 | -0.96175 | 6.05E-07 |
| CCDC11 | CALML6 | -0.97601 | 6.01E-08 |
| BRF1 | CALML6 | 0.957768 | 9.85E-07 |
| GIMAP1 | DEFB106A /// | 0.960782 | 6.84E-07 |
|  | DEFB106B |  |  |
| CCDC65 | DEFB106A /// | 0.951651 | 1.92E-06 |
|  | DEFB106B |  |  |
| RDH10 | DEFB106A /// | 0.968938 | 2.16E-07 |
|  | DEFB106B |  |  |
| VPS18 | WFDC9 | -0.95189 | 1.87E-06 |
| CCDC11 | ZNF485 | 0.968029 | 2.49E-07 |
| BRF1 | ZNF485 | -0.97176 | 1.35E-07 |
| NLRP5 | ZNF485 | -0.95257 | 1.74E-06 |
| CCDC11 | CDH23 | -0.96333 | 4.91E-07 |
| CYP2A6 | DNAJC5G | 0.960958 | 6.69E-07 |
| CCDC11 | DNAJC5G | -0.95102 | 2.04E-06 |
| WDR17 | DNAJC5G | -0.95322 | 1.63E-06 |
| CCDC11 | WDR88 | -0.96017 | 7.38E-07 |
| CATSPER1 | WDR88 | -0.97324 | 1.03E-07 |
| BRF1 | WDR88 | 0.954811 | 1.38E-06 |
| NLRP5 | WDR88 | 0.963964 | 4.50E-07 |
| WDR17 | WDR88 | -0.96062 | 6.98E-07 |
| WDR88 | WDR88 | 0.978602 | 3.41E-08 |
| CYP2A6 | MBD3L2 | -0.96108 | 6.59E-07 |
| CCDC11 | MBD3L2 | 0.971062 | 1.52E-07 |
| CATSPER1 | MBD3L2 | 0.981525 | 1.64E-08 |
| WDR17 | MBD3L2 | 0.965176 | 3.80E-07 |
| ANKAR | MBD3L2 | -0.96971 | 1.91E-07 |
| WDR88 | MBD3L2 | -0.95909 | 8.42E-07 |
| WDR88 | MBD3L2 | -0.96537 | 3.70E-07 |
| PAX8 | ADAMTSL1 | 0.950173 | 2.22E-06 |
| TMEM106A | ADAMTSL1 | -0.96193 | 5.91E-07 |
| ODF4 | MBD3L1 | -0.95259 | 1.74E-06 |
| CCDC11 | MBD3L1 | 0.988363 | 1.65E-09 |
| NLRP5 | MBD3L1 | -0.95047 | 2.16E-06 |
| CCDC11 | FAM46D | 0.950902 | 2.07E-06 |
| ADAM32 | SERPINB11 | -0.96303 | 5.11E-07 |
| NEDD1 | DSCR10 | 0.957306 | 1.04E-06 |
| CYP2A6 | ABHD11 | -0.95662 | 1.13E-06 |
| CATSPER1 | ABHD11 | 0.965901 | 3.43E-07 |
| WDR88 | ABHD11 | -0.96721 | 2.82E-07 |
| WDR88 | ABHD11 | -0.96596 | 3.40E-07 |
| ADAMTSL1 | ABHD11 | 0.953065 | 1.66E-06 |
| MBD3L1 | GAMT | -0.95202 | 1.85E-06 |
| TMEM106A | PTPRC | -0.9591 | 8.41E-07 |
| TMEM106A | RAPH1 | -0.95232 | 1.79E-06 |
| MAN1A2 | RAPH1 | 0.993202 | 1.13E-10 |
| MAN1A2 | SMCR8 | 0.993981 | 6.16E-11 |
| SMCR8 | SMCR8 | 0.998425 | 7.62E-14 |
| PDE7A | LACTB | 0.95717 | 1.06E-06 |
| BNC1 | BNC1 | 0.951039 | 2.04E-06 |
| ODF4 | TAF8 | 0.951929 | 1.86E-06 |
| KLK8 | SLAMF6 | -0.96746 | 2.72E-07 |
| SCARB1 | ZSCAN20 | -0.95303 | 1.66E-06 |
| C4orf33 | RHBDL2 | -0.95553 | 1.27E-06 |
| SERPINB11 | RHBDL2 | 0.951318 | 1.98E-06 |
| ARMCX4 | CRB2 | -0.95373 | 1.55E-06 |
| FAM122C | WBP2NL | -0.97404 | 8.89E-08 |
| TMEM106A | TMEM74 | -0.95533 | 1.30E-06 |
| CATSPER1 | TIGD4 | 0.960038 | 7.50E-07 |
| FAM18B2 | C21orf67 | 0.989391 | 1.04E-09 |
| FAM71A | NLRP11 | 0.966506 | 3.14E-07 |
| CLEC4F | NLRP11 | 0.965087 | 3.85E-07 |
| C21orf67 | ATP6V1C2 | 0.957045 | 1.07E-06 |
| PTPRC | CLDN19 | -0.96369 | 4.68E-07 |
| TIMD4 | KIF6 | 0.952698 | 1.72E-06 |
| DDR1 | TAGAP | 0.95111 | 2.03E-06 |
| PRR22 | TAGAP | 0.956625 | 1.12E-06 |
| HIPK1 | TAGAP | 0.96528 | 3.75E-07 |
| KLHL10 | LETM2 | -0.96726 | 2.80E-07 |
| ESX1 | BSND | 0.965532 | 3.62E-07 |

The authors have commenced performing log transformation oriented normalization and moved further with a primary cue gathering via Principal Component Analysis (PCA). It is also reported that a very few number of overlain genes befall from breast and tumor profiles. Pearson correlation coefficients exhibit stout propensity of breast stromal genes with breast data and prostate stromal genes with prostate data (Figure 1) (Planche et al. 2011). To add to further consolidation of the outcome, survival analysis was carried out using Univariate Cox approach that highlighted genes whose expression levels were crucially associated with the patient survival. The downloaded dataset has no observable missing values in the cells to impute; rather the blank entries are subsidiary to gene and probe ids.

Technically deduced from the background meta-analysis of the subject, we may decipher that cancer will need a host medium (tissue) to proliferate to the other cells/ tissues/ organs. The metastasis front of cancer would seek for the favorable restructuring of the basal tissue framework. From the anticancer therapeutic vantage, hence, it renders incumbent that the oncogenes and stromal response must be equally thrusted.

Through this exemplar multifaceted exegesis, we objectivize to construe the following:

a. Contrivance of differentially expressed genes (DEG)
b. GRN reconstruction, and
c. Decoding functionally coherent genes (eliciting anonymous genes) in accord to iso-form expression.
d. Designing a classifier (machine learning approach) that embraces a threshold value of gene expression that triggers ambient oncological desmoplastic response.

From statistical standpoint, the data concerned is *paired*, i.e. two different conditions (cancerous and normal, here) hybridized on the same slide. A recce exhibits noticeable gene entries that outlie the tightly stratified expression space, as can be derived from the Fig. 3. The dataset dimension of 54675 features tacitly conveys the infestation of multiple gene entries associated with diverse probes. However, a cursory reconnaissance shall also establish that there is only one replicate to each experimental condition.

In recent years, the molecular data has become reverently large. R has evolved as the *defacto* tool for genomic data analysis attributable to its IDE, flexibility and workflow control. Amongst others Python is a viable option too. Biopython is a dedicated version of the language for biological data analytics. However, R has an edge over other languages in terms of packages (functionalities) to cope with the multidimensional data. Being open-source and fully distributable adds to the prowess as well.

## Author Contribution

**Conception and design:** Rajni Jaiswal, Shaurya Jauhari.

**Development of methodology:** Rajni Jaiswal

**Analysis and interpretation of data (e.g., statistical analysis, biostatistics, computational analysis):** Rajni Jaiswal, Shaurya Jauhari.

**Writing, review, and/or revision of the manuscript:** Shaurya Jauhari, S.A.M. Rizvi.

**Study supervision:** S.A.M. Rizvi.

## Stromal Data Analysis: R Script file

~~~
# Installing GEOquery

source(“http://www.bioconductor.org/biocLite.R”)
biocLite(“GEOquery”)

# Loading GEO file with GEOquery

library(Biobase)
library(GEOquery)

#Download GPL file, put it in the current directory, and load it:
gpl570 **<-** getGEO(‘GPL570’, destdir**=**“.”)

#Or, open an existing GPL file:
gpl570 **<-** getGEO(filename**=**‘GPL570.soft’)

# Handpicked description (three columns: ID, Gene Symbol, Gene Title).

Table(gpl570**) [**c(“ID”,”Gene Symbol”,”Gene Title”)]
IDs **<-** attr(dataTable(gpl570), “table”)[, c(“ID”, “Gene Symbol”, “Gene Title”)]

# Extract the expression values from the dataset

# line 64 contains field names

DS_Main **<-** read.table(“GSE26910_series_matrix.txt.gz”, skip **=** 63, header **= TRUE**, sep **=** “\t”, fill **= TRUE)**

# Remove the last line from the matrix that says “!series_matrix_table_end”

DS_Main **<-** DS_Main**[-**54676, ]

# Merging the annotation information to the expression values matrix and rejecting null entries.

names(IDs)[1**] <-** “ID_REF”
DS **<-** merge(IDs,DS_Main, by **=** “ID_REF”)
DS[DS **==** ““**] <-NA**
DS **<-** na.omit(DS)

# Reordering of respective breast cancer and prostate cancer datsets.
# Prostate Normal [1:6], Prostate Tumor [7:12], Breast Normal [13:18], Breast Tumor [19:24]

WorkDS **<-** DS [c(4,6,8,10,12,14, 5,7,9,11,13,15, 16,18,20,22,24,26, 17,19,21,23,25,27)]

# RowMeans calculation

ProstateNormalMean **<-** rowMeans(log2(WorkDS[,1:6]))
ProstateTumorMean **<-** rowMeans(log2(WorkDS[,7:12]))
BreastNormalMean **<-** rowMeans(log2(WorkDS[,13:18]))
BreastTumorMean **<-** rowMeans(log2(WorkDS[,19:24]))

# MA-Plot

par(mfrow**=**c(1,2))
ProstateMean **<-** rowMeans(log2(WorkDS[, 1:12]))
BreastMean **<-** rowMeans(log2(WorkDS[, 13:24]))
plot(ProstateMean, ProstateTumorMean**-**ProstateNormalMean, main**=**“MA Plot for Prostate data”, pch**=**16, cex**=**0.35)
hold()
plot(BreastMean, BreastTumorMean-BreastNormalMean, main="MA Plot for Breast data", pch=16, cex=0.35)

# Rough draft of extreme probes

DS[which.min(BreastTumorMean**-**BreastNormalMean), ] ### most negatively expressed breast gene
DS[which.min(ProstateTumorMean**-**ProstateNormalMean), ] ### most negatively expressed prostate gene
DS[which.max(ProstateTumorMean**-**ProstateNormalMean), ] ### most positively expressed prostate gene
DS[which.max(BreastTumorMean**-**BreastNormalMean), ] ### most positively expressed breast gene

# Standard Deviation calculation for t-test

install.packages(genefilter)
library(genefilter)
ProstateNormalSD **<-** rowSds(log2(WorkDS[,1:6]))
ProstateTumorSD **<-** rowSds(log2(WorkDS[,7:12]))
BreastNormalSD **<-** rowSds(log2(WorkDS[,13:18]))
BreastTumorSD **<-** rowSds(log2(WorkDS[,19:24]))

# t-test calculation and histogram plot

par(mfrow**=**c(1,2))
Prostate_ttest **<-(**ProstateTumorMean**-**ProstateNormalMean**)/**sqrt(ProstateTumorSD**^**2**/**6 **+**
ProstateNormalSD**^**2**/**6)
hist(Prostate_ttest,nclass**=**100)
hold()
Breast_ttest **<- (**BreastTumorMean**-**BreastNormalMean**)/**sqrt(BreastTumorSD**^**2**/**6 **+** BreastNormalSD**^**2**/**6)
hist(Breast_ttest, nclass**=**100)

# p-value calculation and histogram plot

Prostate_pval **<-** 2**\*(**1**-**pt(abs(Prostate_ttest),5))
Breast_pval **<-** 2**\*(**1**-**pt(abs(Breast_ttest),5))
par(mfrow**=**c(1,2))
hist(Prostate_pval, nclass**=**100)
hold()
hist(Breast_pval, nclass **=** 100)

# volcano Plot

par(mfrow**=**c(1,2))
plot(ProstateTumorMean**-**ProstateNormalMean, **-**log10(Prostate_pval), main **=**“Volcano Plot@Prostate tissue”, xlab**=** “Sample Mean Difference”, ylab**=** “-log10(p value)”, pch**=**16, cex**=**0.35)
hold()
plot(BreastTumorMean**-**BreastNormalMean, **-**log10(Breast_pval), main **=**“Volcano Plot@Breast tissue”, xlab**=** “Sample Mean Difference”, ylab**=** “-log10(p value)”, pch**=**16, cex**=**0.35)

# Boxplots for the normal data and its log transformed version.(Log2 transformation applied)

par(mfrow **=** c(1, 2))
boxplot(WorkDS, col **=** c(2,3,2,3,2,3,2,3,2,3,2,3), main **=** “Expression values pre-normalization”,
   xlab **=** “Slides”, ylab **=** “Expression”, las **=** 2, cex.axis **=** 0.7)
hold()
boxplot(log2(WorkDS), col **=** c(2,3,2,3,2,3,2,3,2,3,2,3), main **=** “Expression values post-log-transformation”,
   xlab **=** “Slides”, ylab **=** “Expression”, las **=** 2, cex.axis **=** 0.7)
abline(0, 0, col = "black")

# Check Normality

par(mfrow**=**c(1,2))
qqnorm(Prostate_ttest, main **=** “QQ Plot@Prostate Data”)
qqline(Prostate_ttest)
hold()
qqnorm(Breast_ttest, main **=** “QQ Plot@Breast Data”)
qqline(Breast_ttest)

# Elucidating genes with particular p-values.

**for (**i **in** c(0.01, 0.05, 0.001, 1e**-**04, 1e**-**05, 1e**-**06, 1e**-**07))
print(paste(“genes with p-values smaller than”,i, length(which(Prostate_pval **<** i**))))**
**for (**i **in** c(0.01, 0.05, 0.001, 1e**-**04, 1e**-**05, 1e**-**06, 1e**-**07))
print(paste(“genes with p-values smaller than”,i, length(which(Breast_pval **<** i**))))**

# Plot heatmap of differentially expressed genes: Genes are differentially expressed if its p-value is under a given threshold, which must be smaller than the usual 0.05 or 0.01 due to multiplicity of tests

BreastDEGenes **<-** data.frame(which(Breast_pval **<** 0.01))
ProstateDEGenes **<-** data.frame(which(Prostate_pval **<** 0.01))

ProstateDEGenesData **<-** ProstateDEGenes[, 1]
BreastDEGenesData **<-** BreastDEGenes[, 1]

ProstateData **<-** as.matrix(WorkDS[ProstateDEGenesData, 1:12])
heatmap(ProstateData, col **=** topo.colors(100), cexRow **=** 0.5)

BreastData **<-** as.matrix(WorkDS[BreastDEGenesData, 13:24])
heatmap(BreastData, col **=** topo.colors(100), cexRow **=** 0.5)

# List of differentially expressed genes.

#Breast Data
BDEG **<-** matrix(nrow **=** nrow(BreastDEGenes), ncol **=** 1)
**for(**i **in** 1:nrow(BreastDEGenes))
BDEG[i,**]<-** paste(DS[BreastDEGenes[i,], “ID_REF”]) BDEG **<-** as.data.frame(BDEG)
names(BDEG)[1**] <-** “ID_REF”
FinalBDEG **<-** merge(BDEG,DS)
BDEG **<-** merge(BDEG, IDs, by **=** ‘ID_REF’)
view(BDEG)

#Prostate Data
PDEG **<-** matrix(nrow **=** nrow(ProstateDEGenes),ncol **=** 1)
**for(**i **in** 1:nrow(ProstateDEGenes)) PDEG[i,**] <-** paste(DS[ProstateDEGenes[i,], “ID_REF”])
PDEG **<-** as.data.frame(PDEG)
names(PDEG)[1**] <-** “ID_REF”
FinalPDEG **<-** merge(PDEG,DS)
PDEG **<-** merge(PDEG, IDs, by **=** ‘ID_REF’)
view(PDEG)

##Intersecting transcripts in breast and prostate cancer types as marked in the dataset

BDEG**$**match **<-** match(BDEG**$**location, PDEG**$**location, nomatch**=**0)

# Reordering of respective breast cancer and prostate cancer datsets.

# Prostate Normal [1:6], Prostate Tumor [7:12], Breast Normal [13:18], Breast Tumor [19:24]

FinalPDEG **<-** FinalPDEG [c(4,6,8,10,12,14, 5,7,9,11,13,15, 16,18,20,22,24,26, 17,19,21,23,25,27)]
WorkFinalPDEG **<-** FinalPDEG[1:12]

FinalBDEG **<-** FinalBDEG [c(4,6,8,10,12,14, 5,7,9,11,13,15, 16,18,20,22,24,26, 17,19,21,23,25,27)]
WorkFinalBDEG **<-** FinalBDEG[13:24]

##Prostate data regrerssion analysis(linear model)

par(mfrow**=**c(1,6))
plot(log2(WorkFinalPDEG**$**GSM662756),log2(WorkFinalPDEG**$**GSM662757), pch **=** 16, cex **=** 1.3, col **=**
c(“blue”,”red”))
abline(lm(log2(WorkFinalPDEG**$**GSM662756**) ∼** log2(WorkFinalPDEG**$**GSM662757)), col**=** 1) plot(log2(WorkFinalPDEG**$**GSM662758),log2(WorkFinalPDEG**$**GSM662759), pch **=** 16, cex **=** 1.3, col **=** c(“blue”,”red”))
abline(lm(log2(WorkFinalPDEG**$**GSM662758**) ∼** log2(WorkFinalPDEG**$**GSM662759)), col**=** 1)
plot(log2(WorkFinalPDEG**$**GSM662760),log2(WorkFinalPDEG**$**GSM662761), pch **=** 16, cex **=** 1.3, col **=**
c(“blue”,”red”))
abline(lm(log2(WorkFinalPDEG**$**GSM662760**) ∼** log2(WorkFinalPDEG**$**GSM662761)), col**=** 1) plot(log2(WorkFinalPDEG**$**GSM662762),log2(WorkFinalPDEG**$**GSM662763), pch **=** 16, cex **=** 1.3, col **=** c(“blue”,”red”))
abline(lm(log2(WorkFinalPDEG**$**GSM662762**) ∼** log2(WorkFinalPDEG**$**GSM662763)), col**=** 1) plot(log2(WorkFinalPDEG**$**GSM662764),log2(WorkFinalPDEG**$**GSM662765), pch **=** 16, cex **=** 1.3, col **=** c(“blue”,”red”))
abline(lm(log2(WorkFinalPDEG**$**GSM662764**) ∼** log2(WorkFinalPDEG**$**GSM662765)), col**=** 1)
plot(log2(WorkFinalPDEG**$**GSM662766),log2(WorkFinalPDEG**$**GSM662767), pch **=** 16, cex **=** 1.3, col **=** c(“blue”,”red”))
abline(lm(log2(WorkFinalPDEG**$**GSM662766**) ∼** log2(WorkFinalPDEG**$**GSM662767)), col**=** 1)

##Gene Set Enrichment Analysis

library(genefilter)
library(GSEABase)
Breast_GSEA **<-** GeneSetCollection(WorkFinalBDEG, setType **=** KEGGCollection())
Prostate_GSEA **<-** GeneSetCollection(WorkFinalPDEG, setType **=** KEGGCollection())

##Correlation Analysis

##Breast

WorkFinalBDEG **<-** read.csv(“WorkFinalBDEG_GSEAFiltered.csv”) ## Import filtered annotation file from MeV.
btemp **<-** WorkFinalBDEG
btemp**$**ID_REF **<-NULL**
btemp **<-** log2(btemp)
pairs(btemp)
BreastCorrelationMatrix **<-** cor(t(as.matrix(btemp)))
BreastCorMat **<-** as.data.frame(BreastCorrelationMatrix)
rownames(BreastCorMat**) <-** WorkFinalBDEG**$**ID_REF
colnames(BreastCorMat**) <-** WorkFinalBDEG**$**ID_REF

##Prostate

WorkFinalPDEG **<-** read.csv(“WorkFinalPDEG_GSEAFiltered.csv”) ## Import filtered annotation file from MeV.
ptemp **<-** WorkFinalPDEG
ptemp**$**ID_REF **<-NULL**
ptemp **<-** log2(ptemp)
pairs(ptemp)
ProstateCorrelationMatrix **<-** cor(t(as.matrix(ptemp)))
ProCorMat**<-** as.data.frame(ProstateCorrelationMatrix)
rownames(ProCorMat**) <-** WorkFinalPDEG**$**ID_REF
colnames(ProCorMat**)<-** WorkFinalPDEG**$**ID_REF

### Feature Selection: Clustering of robustly entwined genes.

install.packages(“gplots”)
install.packages(“Hmisc”)
library(Hmisc)
library(gplots)
heatmap.2(ProstateCorrelationMatrix, main**=**“Hierarchical Cluster”,
dendrogram**=**“column”,trace**=**“none”,col**=**greenred(10))
heatmap.2(1**-**abs(ProstateCorrelationMatrix), distfun**=**as.dist, trace**=**“none”) heatmap.2(BreastCorrelationMatrix, main**=**“Hierarchical Cluster”,
dendrogram**=**“column”,trace**=**“none”,col**=**greenred(10))
heatmap.2(1**-**abs(BreastCorrelationMatrix), distfun**=**as.dist, trace**=**“none”)

##Prostate Data

library(caret)
HighlyCorrelated **<-** findCorrelation(ProstateCorrelationMatrix, cutoff **=** 0.95, verbose **= TRUE**, names **= FALSE)**
print(HighlyCorrelated)
WorkFinalPDEG[HighlyCorrelated,1]
IDs[WorkFinalPDEG[HighlyCorrelated,1],c(2,3)]
**for(**i **in** 2:nrow(BreastCorMat))
 **{**
  **for(**j **in** 1:ncol(BreastCorMat**)-**1)
   **{**
     **if(**i**>**j)
     **{**
     out **<-** c (rownames(BreastCorMat[i,]), colnames(BreastCorMat[j]), BreastCorMat[i,j]) write.table(out, file**=**“output.txt”, append**=TRUE**, sep**=** “ “)
     **}**
    **else**
   **break**
  **}**
**}**

#Network Ready Matrix Format (Function) // Credit: <u>http://www.sthda.com</u>

flattenCorrMatrix **<-function(**cmat, pmat**) {**
  ut **<-** upper.tri(cmat)
  data.frame(
   row **=** rownames(cmat)[row(cmat)[ut]],
   column **=** rownames(cmat)[col(cmat)[ut]],
   cor **=(**cmat)[ut],
   p **=** pmat[ut]
  )
**}**

library(Hmisc)

btemp **<-** as.matrix(btemp)
rownames(btemp**)<-** WorkFinalBDEG**$**ID_REF
BreastNet **<-**rcorr(t(btemp))
BreastNetworkInputMatrix**<-** flattenCorrMatrix(BreastNet**$**r, BreastNet**$**P)

#lets map the gene names to row and column entries
BreastNetworkInputMatrix**$**row **<-**IDs[WorkFinalBDEG[BreastNetworkInputMatrix**$**row,1],2]
BreastNetworkInputMatrix**$**column **<-**IDs[WorkFinalBDEG[BreastNetworkInputMatrix**$**column,1],2]
ptemp **<-** as.matrix(ptemp)
rownames(ptemp**)<-** WorkFinalPDEG**$**ID_REF
ProstateNet **<-**rcorr(t(ptemp))
ProstateNetworkInputMatrix **<-** flattenCorrMatrix(ProstateNet**$**r, ProstateNet**$**P)
ProstateNetworkInputMatrix**$**row **<-**IDs[WorkFinalPDEG[ProstateNetworkInputMatrix**$**row,1],2]
ProstateNetworkInputMatrix**$**column **<-**IDs[WorkFinalPDEG[ProstateNetworkInputMatrix**$**column,1],2]

symnum(BreastCorrelationMatrix)
symnum(ProstateCorrelationMatrix)

install.packages(“corrplot”)
library(corrplot)
corrplot(BreastCorrelationMatrix, type**=**“upper”, order**=**“hclust”, tl.col**=**“black”, tl.srt**=**45)
corrplot(ProstateCorrelationMatrix, type**=**“upper”, order**=**“hclust”, tl.col**=**“black”, tl.srt**=**45)

install.packages(“PerformanceAnalytics”)
library(PerformanceAnalytics)
chart.Correlation(BreastCorrelationMatrix, histogram**= TRUE**, pch**=** 19)
chart.Correlation(ProstateCorrelationMatrix, histogram**= TRUE**, pch**=** 19)

col**<-** colorRampPalette(c(“blue”, “white”, “red”))(20)
heatmap(x **=** BreastCorrelationMatrix, col **=** col, symm **= TRUE)**
heatmap(x **=** ProstateCorrelationMatrix, col **=** col, symm **= TRUE)**

## top candidates which manifest low p-value and high correlation.
BreastFinal **<-** BreastNetworkInputMatrix[which(abs(BreastNetworkInputMatrix**$**cor**) >** 0.95 **|**
BreastNetworkInputMatrix**$**p **<** 0.0000001), c(1,2,3,4)]
ProstateFinal **<-** ProstateNetworkInputMatrix[which(abs(ProstateNetworkInputMatrix**$**cor**) >** 0.95 **|**
ProstateNetworkInputMatrix**$**p **<** 0.0000001), c(1,2,3,4)]
write.csv(BreastFinal, “BreastFinalTest.csv”)
write.csv(ProstateFinal, “ProstateFinalTest.csv”)

##Intersecting transcripts in breast and prostate cancer types as marked in the dataset

BDEG**$**match **<-** match(BDEG**$**location, PDEG**$**location, nomatch**=**0)

# Reordering of respective breast cancer and prostate cancer datsets.

# Prostate Normal [1:6], Prostate Tumor [7:12], Breast Normal [13:18], Breast Tumor [19:24]

FinalPDEG **<-** FinalPDEG [c(4,6,8,10,12,14, 5,7,9,11,13,15, 16,18,20,22,24,26, 17,19,21,23,25,27)]
WorkFinalPDEG **<-** FinalPDEG[1:12]

FinalBDEG **<-** FinalBDEG [c(4,6,8,10,12,14, 5,7,9,11,13,15, 16,18,20,22,24,26, 17,19,21,23,25,27)]
WorkFinalBDEG **<-** FinalBDEG[13:24]

##Prostate data regrerssion analysis(linear model)

par(mfrow**=**c(1,6))
plot(log2(WorkFinalPDEG**$**GSM662756),log2(WorkFinalPDEG**$**GSM662757), pch **=** 16, cex **=** 1.3, col **=** c(“blue”,”red”))
abline(lm(log2(WorkFinalPDEG**$**GSM662756**) ∼** log2(WorkFinalPDEG**$**GSM662757)), col**=** 1)
plot(log2(WorkFinalPDEG**$**GSM662758),log2(WorkFinalPDEG**$**GSM662759), pch **=** 16, cex **=** 1.3, col **=** c(“blue”,”red”))
abline(lm(log2(WorkFinalPDEG**$**GSM662758**) ∼** log2(WorkFinalPDEG**$**GSM662759)), col**=** 1)
plot(log2(WorkFinalPDEG**$**GSM662760),log2(WorkFinalPDEG**$**GSM662761), pch **=** 16, cex **=** 1.3, col **=** c(“blue”,”red”))
abline(lm(log2(WorkFinalPDEG**$**GSM662760**) ∼** log2(WorkFinalPDEG**$**GSM662761)), col**=** 1)
plot(log2(WorkFinalPDEG**$**GSM662762),log2(WorkFinalPDEG**$**GSM662763), pch **=** 16, cex **=** 1.3, col **=** c(“blue”,”red”))
abline(lm(log2(WorkFinalPDEG**$**GSM662762**) ∼** log2(WorkFinalPDEG**$**GSM662763)), col**=** 1)
plot(log2(WorkFinalPDEG**$**GSM662764),log2(WorkFinalPDEG**$**GSM662765), pch **=** 16, cex **=** 1.3, col **=** c(“blue”,”red”))
abline(lm(log2(WorkFinalPDEG**$**GSM662764**) ∼** log2(WorkFinalPDEG**$**GSM662765)), col**=** 1)
plot(log2(WorkFinalPDEG**$**GSM662766),log2(WorkFinalPDEG**$**GSM662767), pch **=** 16, cex **=** 1.3, col **=** c(“blue”,”red”))
abline(lm(log2(WorkFinalPDEG**$**GSM662766**) ∼** log2(WorkFinalPDEG**$**GSM662767)), col**=** 1)

##Gene Set Enrichment Analysis

library(genefilter)
library(GSEABase)
Breast_GSEA **<-** GeneSetCollection(WorkFinalBDEG, setType **=** KEGGCollection())
Prostate_GSEA **<-** GeneSetCollection(WorkFinalPDEG, setType **=** KEGGCollection())

# Support Vector Machine Implementation

## Prostate
install.packages(“e1071”)
library(e1071)
temp1 **<-** WorkFinalPDEG temp1**$**ID_REF **<-NULL**
temp1 **<-** log2(temp1)
temp1 **<-** t(temp1)
ClassLabels1 **<-** c(rep(1,6),rep**(-**1,6))
DataFrame1 **<-** data.frame(Gene**=**temp1,ClassLabels**=**as.factor(ClassLabels1))
SVMModel1 **<-** svm(ClassLabels1**∼**., data**=**DataFrame1, kernel**=**“linear”, cost**=**10, scale **= FALSE)**
GeneWeights1**<-**t(SVMModel1**$**coefs)%*%SVMModel1**$**SV
sort.list(GeneWeights1) ## Genes 212 and 129 have highest and second highest weights, respectively.
plot(SVMModel1,DataFrame1, Gene.212 **∼** Gene.129)

##Breast

temp2 **<-** WorkFinalBDEG
temp2**$**ID_REF **<-NULL**
temp2 **<-** log2(temp2)
temp2 **<-** t(temp2)
ClassLabels2**<-** c(rep(1,6),rep**(-**1,6))
DataFrame2 **<-** data.frame(Gene**=**temp2,ClassLabels**=**as.factor(ClassLabels2))
SVMModel2 **<-** svm(ClassLabels2**∼**., data**=**DataFrame2, kernel**=**“linear”, cost**=**10, scale **= FALSE)**
GeneWeights2**<-**t(SVMModel2**$**coefs)%*%SVMModel2**$**SV
sort.list(GeneWeights2) ## Genes 346 and 133 have highest and second highest weights, respectively. plot(SVMModel2,DataFrame2, Gene.346 **∼** Gene.133)
install.packages(“kernlab”)
library(kernlab)
x **<-** as.matrix(temp1)
y **<-** matrix(c(rep(1,6),rep**(-**1,6)))
svp **<-** ksvm(x,y,type**=**“C-svc”, prob.model**= TRUE)**
predict (svp, x, type**=** “probabilities”)
~~~

